# Novelty is not Surprise: Human exploratory and adaptive behavior in sequential decision-making

**DOI:** 10.1101/2020.09.24.311084

**Authors:** He A. Xu, Alireza Modirshanechi, Marco P. Lehmann, Wulfram Gerstner, Michael H. Herzog

## Abstract

Classic reinforcement learning (RL) theories cannot explain human behavior in response to changes in the environment or in the absence of external reward. Here, we design a deep sequential decision-making paradigm with sparse reward and abrupt environmental changes. To explain the behavior of human participants in these environments, we show that RL theories need to include surprise and novelty, each with a distinct role. While novelty drives exploration before the first encounter of a reward, surprise increases the rate of learning of a world-model as well as of model-free action-values. Even though the world-model is available for model-based RL, we find that human decisions are dominated by model-free action choices. The world-model is only marginally used for planning but is important to detect surprising events. Our theory predicts human action choices with high probability and allows us to dissociate surprise, novelty, and reward in EEG signals.

## Introduction

Humans seek not only explicit rewards such as money or praise [1–8], but also novelty [9, 10], an intrinsic reward-like signal which is linked to curiosity [9–13]. In the theory of reinforcement learning, novelty is considered as a drive for exploration [11, 14–16], and novelty-driven exploratory actions have been interpreted as steps towards building a model of the world (‘world-model’) which is then used for action planning [17]. A world-model represents implicit knowledge that links actions to observations, such as ‘if I open the door to my kitchen, I will see my fridge’.

However, since the world is much more complex than any model of it, there will occasionally be a mismatch between the expectations arising from the model and the actual observation, e.g., when you return from work and the location of the fridge is suddenly empty because your room-mate has sent it off for repair. Such mismatches generate the feeling of surprise, known to manifest in pupil dilation [18] and EEG signals [19–21]. Whereas the reward prediction error (RPE) is a mismatch between the expected reward and the actual reward, surprise is a mismatch between an expected observation and an actual observation. Behavioral experiments [18, 22–25] and theories [25–28] suggest that surprise helps humans to adapt their behavior quickly to changes in the environment, potentially by modulating synaptic plasticity [29–31].

Surprise is fundamentally different from novelty; if you already know that your fridge would be fetched for repairing, the new arrangement of the kitchen without the fridge is novel but not surprising. However, although there is some agreement that novelty and surprise are two separate notions, it has been debated how they can be formally distinguished [32–34], whether they manifest themselves differently in EEG signals [20, 21, 35, 36], and how they influence learning and decisionmaking [9–13, 18, 22–24, 37].

In this study, we address three questions: First, how do surprise and novelty influence reinforcement learning? Second, what is their relative contribution to exploratory and adaptive behavior? And third, can surprise be distinguished from novelty in human behavioral choices and event related potentials (ERP) of the electroencephalogram (EEG)? We show, via a specifically designed deep sequential decision task and a novel hybrid reinforcement learning model, that we can dissociate contributions of surprise from those of novelty and reward in human behavior and ERP.

Our key findings can be summarized in three points: (i) We find that novelty-seeking explains participants’ exploratory behavior better than alternative exploration strategies such as seeking surprise or uncertainty [38, 39]; (ii) we observe that participants use their world-model only rarely for action planning and mainly to extract moments of surprise; and importantly, (iii) we show that surprise calculated by the world-model does not only modulate the learning of the world-model [22–24, 27] but also the learning of model-free action-values. In particular, we show that such a modulation is necessary to explain participants’ adaptive behavior.

## Results

### Experimental paradigm and human behavior

In order to distinguish between novelty, surprise, and reward, and to study their effects on exploratory and adaptive behavior, we designed an environment (cf.[40]) consisting of 10 states with 4 possible actions per state plus one goal state (Fig. 1A and Fig. 1B). In the human experiments, states were represented as images on a computer screen and actions as four grey disks below the image. Before the experiment, 12 participants were shown all images of the states and were informed that their task was to find the shortest path to the goal image. Throughout the experiment, at each state, participants chose an action (by clicking on one of the grey disks) which brought them to the next image, where they then chose the next action, and so on (Fig. 1A). The episode ended when the goal image was found.

**Figure 1:**
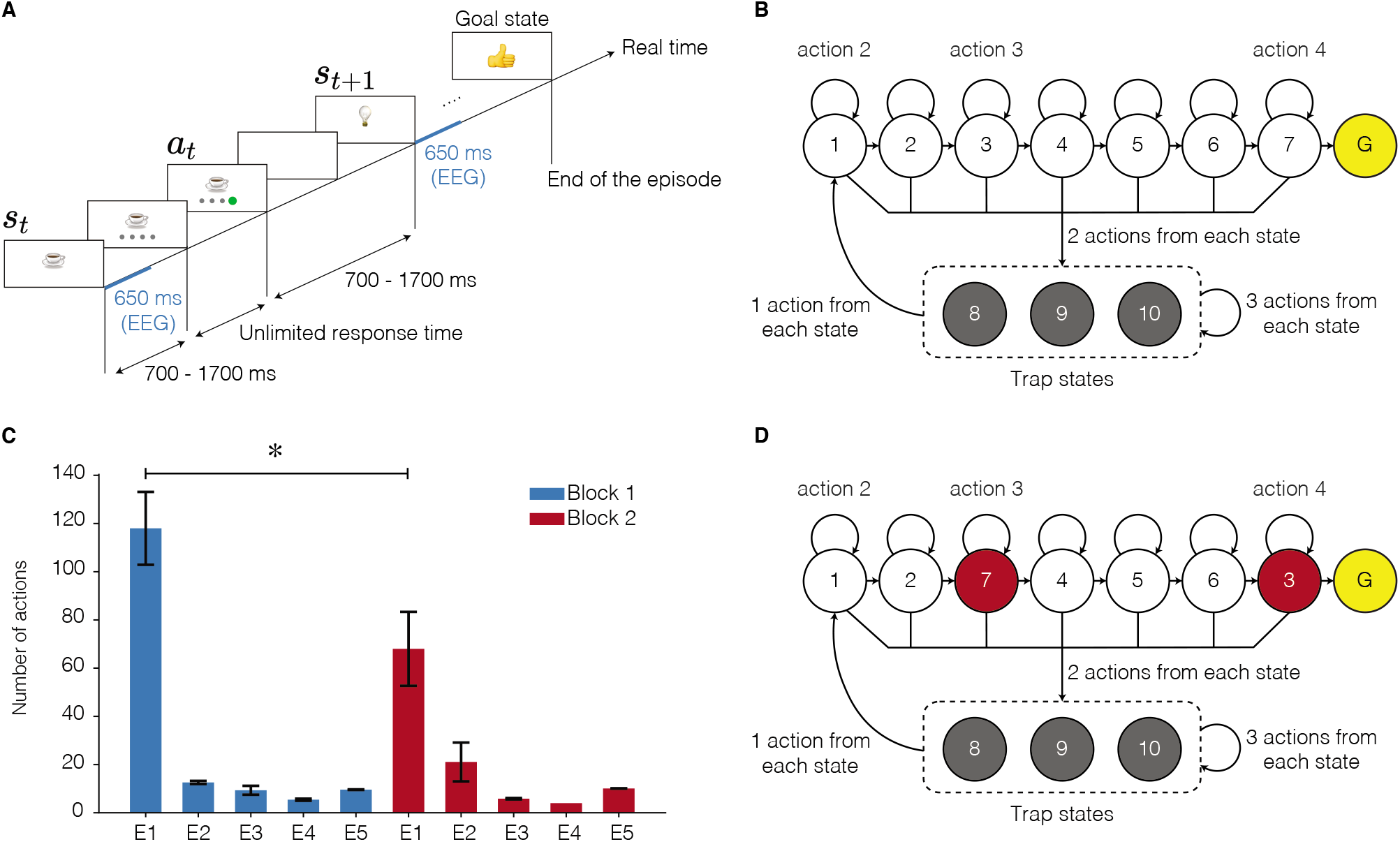
Experimental paradigm. **A.** After image onset, participants had to wait for 700-1700ms (randomly chosen) until grey disks were presented at the bottom of the image. After clicking on one disk, a blank screen was presented for another random interval of 700 to 1700ms. The next image appeared afterwards. Different participants saw different images, but the underlying structure was identical for all participants. The goal image is a ‘thumb-up’ image in this example. The blue lines indicate the window of EEG analysis. **B.** Structure of the environment during block 1. There were 10 states with 4 actions each plus a goal state (G). States 1-7 are *progressing states* and states 8-10 are *trap states.* For each progressing state, one action led participants to the next progressing state, two actions led participants to one of the trap states, and one action made participants stay at the current state. The action which made participants stay at the current state is shown for states 1, 3, and 7, as an example. For each trap state, three actions led participants to one of the trap states, and one action led participants to state 1. Not all action arrows are drawn for the trap states to simplify illustration. **C.** Average number of actions of participants during block 1 (blue) and block 2 (red): The 1st episode of block 2 was significantly shorter than the 1st episode of block 1 (one-sample t-test, p-value=0.035). Error bars show the standard error of the mean. **D.** Environment used in block 2: The images presenting state 3 and state 7 (in red) were swapped. Other transitions remained unchanged.

Unknown to the participants, the non-goal states could be classified into progressing states (1 to 7 in Fig. 1B) and trap states (8 to 10 in Fig. 1B). In Fig. 1B, we arrange progressing states as a path toward the goal. At each progressive state, one action brought participants either to another progressive state closer to the goal or directly to the goal state, two actions brought them to one of the trap states, and one action made them stay at the current state. At each trap state, three actions brought participants to either the same or another trap state, and one action brought them to state 1, at the beginning of the path of progressing states. Note that the underlying structure of the environment, the assignment of images to specific states, and the assignment of action buttons to specific transitions were unknown to the participants.

The experiment was organized in 10 episodes which, unknown to the participants, were divided into two blocks of 5 episodes each. During the 1st episode of the 1st block, participants took between 34 and 214 actions (mean 117 and std 54) until they arrived at the goal (Fig. 1C). They then continued for another 4 episodes, each time starting in a new initial state that had been randomly chosen but kept fixed across participants. After the first episode, participants learned to reach the goal in less than 20 steps (Fig. 1C). After the end of the 5th episode, two states (state 3 and 7 in Fig. 1D) were swapped, without announcing it to the participants. Participants continued for another 5 episodes with the novel layout of the environment (2nd block, Fig. 1D).

In the 1st episode of block 1, participants explored the environment to find the goal, but they received no intermediate reward or other sign of progress while doing this. In our complex environment, if participants followed a purely random exploration strategy (i.e., choosing each action with 1/4 probability), it would take them on average about 10^4^ actions to find the goal, starting at any non-goal state (see Supplementary Material). Our results, suggest that participants followed a non-random strategy for finding the goal (Fig. 1C). Here, we ask whether novelty of states played a role in the way participants chose their actions.

In the 1st episode of block 2, when states 3 and 7 had been swapped, participants were significantly faster in finding the goal state than in the 1st episode of block 1 (Fig. 1C), indicating that they adapted their behavior to the new situation while exploiting knowledge they had acquired before. This observation suggests that surprise triggered by unexpected transitions helped participants to rapidly adapt their behavior.

### Defining Novelty and Surprise

We assume that states that are encountered frequently are experienced by participants as less novel than those encountered for the first time. We, therefore, define the novelty of a state *s* at time *t* as a decreasing function of the number 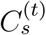 of encounters of state *s* until time *t*. More precisely, we define the novelty of state *s* at time *t* as

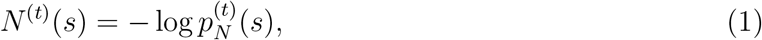

where

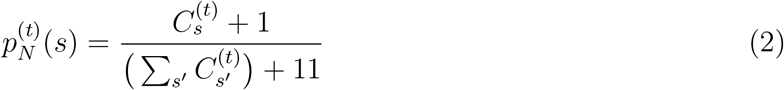

has two different interpretations. First, 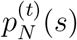 can be seen as the probability of observing state *s* at time *t,* estimated in a Bayesian framework (see Supplementary Material); measures similar to 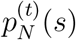, sometimes called ‘density models’, have been used in machine learning community [15]. In the Bayesian interpretation, the numbers 1 in the nominator and 11 in the denominator (11 is the total number of states in the environment) correspond to a uniform prior that makes all states equally likely at time *t* = 0. In the second interpretation, 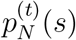 is seen as the empirical frequency of observing state *s_t_* until time *t*. In fact, because one of the counters *C_s’_* increases by one at each time step, the time can be expressed as 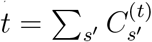. In this interpretation, the numbers 1 in the nominator and 11 in the denominator correspond to the one encounter of each state before the start of the experiment.

With our definition of novelty, at the beginning of the 1st episode in block 1, all states have identical novelty. Since participants often fall into one of the trap states, the novelty of trap states decreases rapidly (Fig. 2A). Hence, before the end of the 1st episode, the novelty is highest for states in the proximity of the goal (Fig. 2B). This observation suggests that seeking novel states will, in our environment, effectively lead a participant closer to the goal, even *before* the participant knows where the goal is located, i.e., before encountering the goal for the first time. When formalizing our hypothesis below, we will use this insight to assign novelty values to states, where novelty values influence behavioral decisions similar to reward-based values.

**Figure 2:**
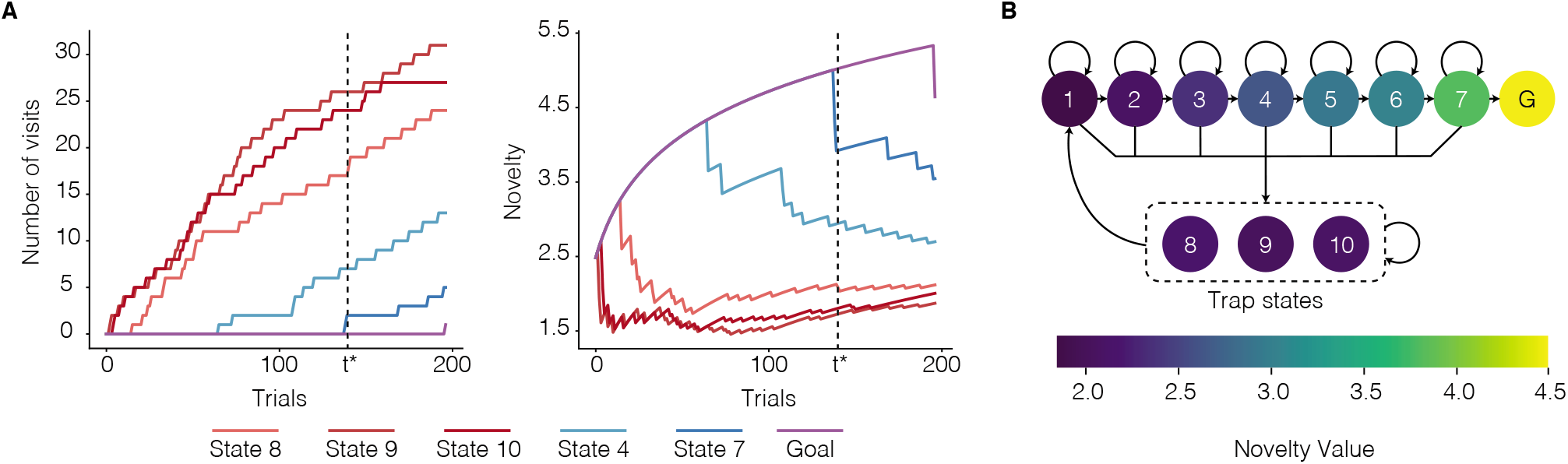
Novelty in episode 1 of block 1. **A.** The number of state visits (left panel) and novelty (right panel) as a function of time for one representative participant: The number of visits increases rapidly for the trap states and remains 0 for a long time for the states closer to the goal. Novelty of each state is defined as the negative log-probability of observing that state (see Eq. 1 and Eq. 2) and, hence, increases for states which are not observed as time passes. The first time participants encounter state 7 (the state before the goal state) is denoted by *t*\*. **B.** Average (over participants) novelty (color coded) at *t*\*: Novelty of each state is a decreasing function of its distance from the goal state.

To navigate in such a complex environment, we assume that participants build an internal model of the environment (‘world-model’), i.e., we hypothesize that participants estimate the probabilities 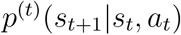 of transitions from a given state *s_t_* to another state *s*_*t*+1_ when performing action *a_t_*. With this assumption, we define surprise as a measure quantifying how ‘unexpected’ the next image (state *s*_*t*+1_) is given the previous state *s_t_* and the chosen action *a_t_*. More precisely, we use a recent measure of surprise motivated by a Bayesian framework for learning in volatile environments, called the ‘Bayes Factor’ surprise [27]. The Bayes Factor surprise of the transition from state *s_t_* to state *s*_*t*+1_ after taking action *a_t_* is

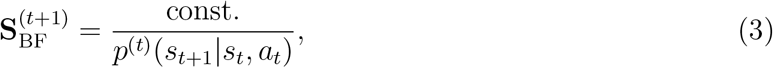

where 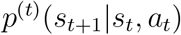 is the conditional probability of observing state *s*_*t*+1_ at time *t* + 1 derived from the present world-model. More precisely, we assume that the world model counts transitions from state *s* to *s*’ under action *a* using either a leaky [21, 41, 42] or a surprise modulated [26, 27] counting procedure, described by the pseudo-count 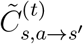. The conditional probability is then

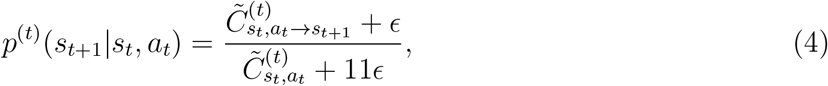

where *ϵ* is a parameter corresponding to a prior in the Bayesian framework, and 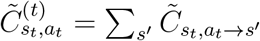 is the pseudo-count of taking action *a_t_* at state *s_t_* (see Supplementary Material). If there is no forgetting or surprise modulation, pseudo-counts are equal to the real counts. Our surprise measure is an increasing function of the state prediction error [5] and Shannon surprise [42, 43] (see Methods) and takes high values during the 1st episode of block 2 whenever participants encounter states 3 or 7 or transit from state 3 or 7 to another state (Fig. 3A and Fig. 3B).

**Figure 3:**
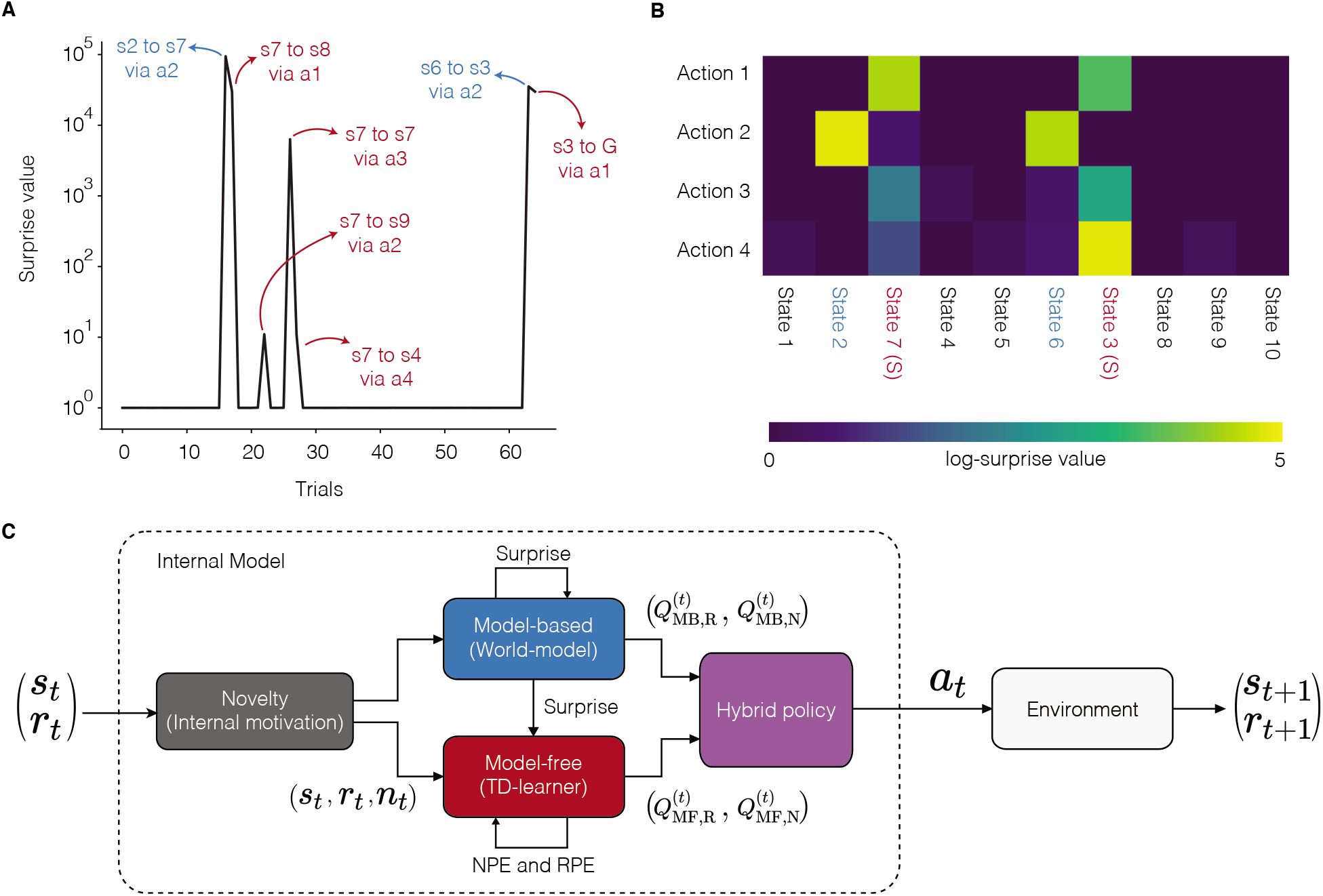
Surprise as a modulator of the learning rate in episode 1 of block 2. **A.** Surprise as a function of time since the start of block 2 for one representative participant: Surprise values are almost zero most of the time, because the participant has already learned the transitions in the environment during block 1. The surprising transitions are the ones to the swapped states (blue) and the ones from the swapped states (red). **B.** Maximal log-surprise values (yellow=large surprise) during the 1st episode of block 2, averaged over all participants. The swapped states are marked in red and the states before them in blue. One action from each swapped state is not surprising, i.e., the action leading participants to trap states both before and after the swap. **C.** Block diagram of the SurNoR algorithm: Information of state *s_t_* and reward *r_t_* at time *t* is combined with novelty *n_t_* (grey block) and passed on to the world-model (blue block, implementing the model-based branch of SurNor) and TD learner (red block, implementing the model-free branch). The surprise value computed by the world-model modulates the learning rate of both the TD-learner and the world-model. The output of each block is a pair of Q-values i.e, Q-values for estimated reward *Q*_MF,R_ and *Q*_MB,R_ as well as for estimated novelty *Q*_MF,N_ and *Q*_MB,N_. The hybrid policy (in purple) combines these values.

### The SurNoR algorithm: Distinct contributions of novelty and surprise to behavior

We hypothesize that participants use novelty to explore the environment and surprise to modulate the rate of learning. The hypothesis is formalized in the form of the Surprise-Novelty-Reward (SurNoR) algorithm and tested given the behavioral data of 12 participants.

Novelty in SurNoR plays a role analogous to that of reward. For example, in standard Temporal-Difference (TD) Learning, a reward-based Q-value *Q_R_*(*s,a*) is associated with each state-action pair (*s,a*) [17]. The Q-value *Q_R_*(*s,a*) estimates the mean discounted reward that can be collected under the current policy when starting from state *s* and action *α*. The reward prediction error RPE, derived from *Q_R_*(*s,a*), serves as a learning signal even for states a few steps away from the goal [17]. Analogously, in the SurNoR model, novelty is a reward-like signal with associated novelty-based Q-values *Q_N_*(*s,a*) and an associated novelty prediction error (NPE) derived from *Q_N_*(*s,α*). In the SurNoR model, the two sets of Q-values, reward-based and novelty-based, are used in a hybrid model [5, 6] that flexibly combines model-based with model-free action selection policies (Fig. 3C).

Surprise in SurNoR is derived from a mismatch between observations of the next state and predictions arising from the world-model embedded in the model-based branch of SurNoR. To adapt both model-based and model-free policies of the SurNoR algorithm, surprise is used in two different ways. First, high values of surprise systematically lead to a larger learning rate for the update of the world-model than smaller ones, consistent with earlier models [25, 27]. Second, going beyond previous models of behavior [18, 22–24, 28], surprise also influences the learning rate of the model-free reinforcement learning branch.

We predict that, if the behavior of participants is well described by the SurNoR algorithm, they should use an action policy that attracts them to novel states, in particular during the 1st episode of block 1. If participants do not exploit novelty, standard (potentially hybrid) reinforcement learning schemes in combination with one of several alternative exploration strategies (see next section) should be sufficient to explain the behavior. Furthermore, we predict that, if the behavior of participants is well described by the SurNoR algorithm, then surprising events during the 1st episode of block 2 should significantly change the behavior of participants; if participants do not exploit surprise, standard hybrid models combining model-based and model-free reinforcement learning [5, 6] should be sufficient to describe the behavior.

### Both surprise and novelty are needed to explain behavior

To test our hypothesis, and to test whether all elements of SurNoR (i.e., surprise, novelty, and hybrid policy) are necessary for explaining behavior or whether a simpler or an alternative model would have the same explanatory power, we compared 12 alternative algorithms with SurNoR: 4 model-based (MB), 4 model-free (MF), and 3 hybrid (Hyb) algorithms plus a null algorithm based on a random choice (RC) of actions (Fig. 4A). Seven out of these 12 algorithms use surprise (+S), and five use novelty (+N, see Supplementary Material). As an alternative for surprise modulation, we used a constant learning rate [20, 21, 41, 42] (all algorithms without +S), and as alternative exploration strategies, we used optimistic initialization (+OI) [17] and uncertainty (surprise) seeking [38, 39] (+U); see below for more explanations and Supplementary Material for details.

**Figure 4:**
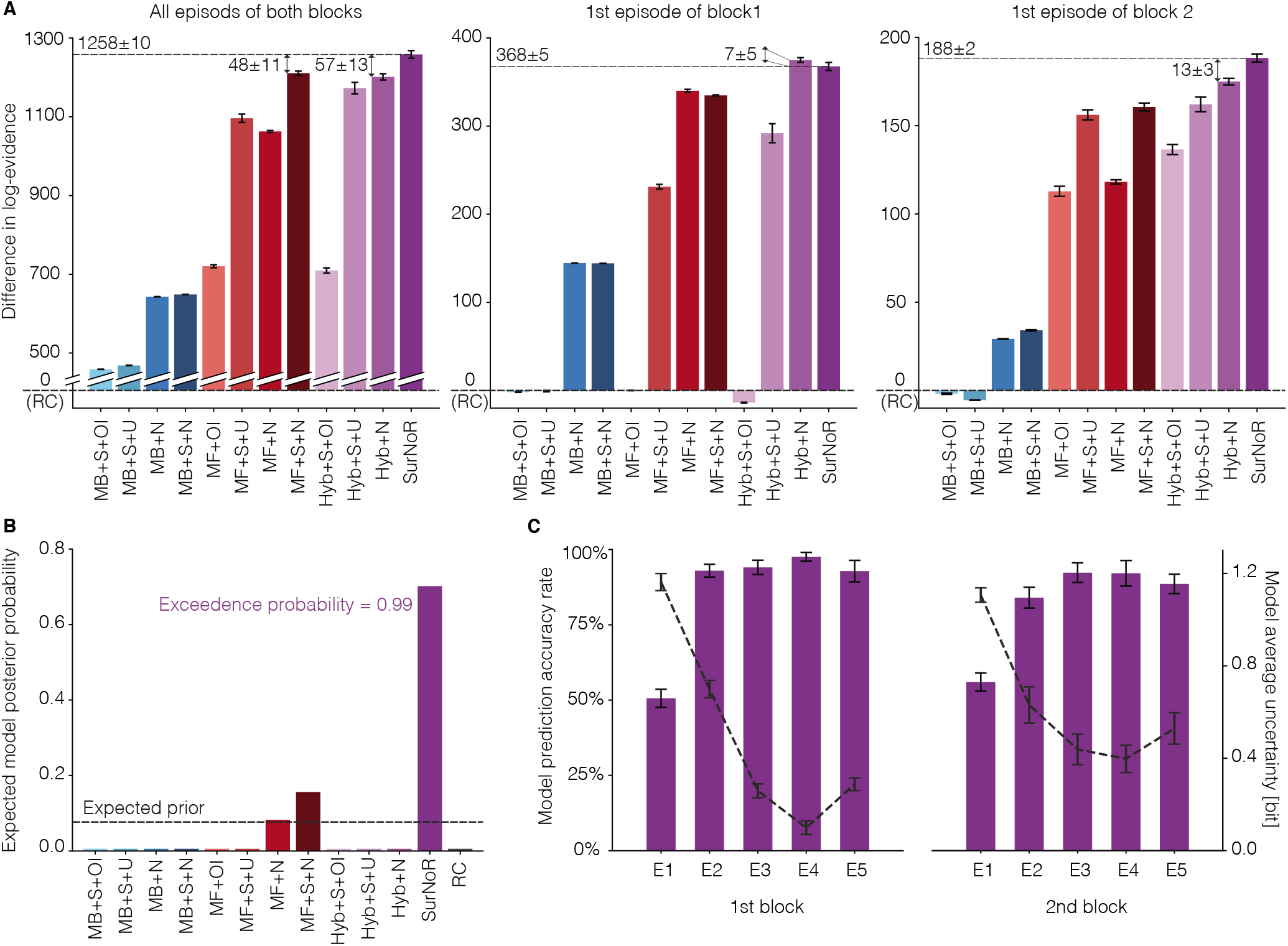
Model comparison of model-based (MB, blue bars), model-free (MF, red bars), and hybrid algorithms (Hyb and SurNoR, purple bars). Exploratory behavior is either induced by optimistic initialization (+OI), uncertainty-seeking (+U), unbiased random action choices (RC), or novelty-seeking (+N); e.g., a model-based algorithm with novelty seeking is denoted as MB+N. SurNoR and the model-free or hybrid algorithms annotated with ‘+S’ use surprise to modulate the learning rate of the model-free TD learner; SurNoR and all algorithms annotated with ‘+S’ use surprise modulation also during model building (see Methods). **A.** Difference in log-evidence (with respect to RC) for the algorithms for all episodes of both blocks (left panel), the 1st episode of block 1 (middle), and the 1st episode of block 2 (right panel). High values indicate good performance; differences greater than 3 or 10 are considered as significant or strongly significant, respectively (see Methods); a value of 0 corresponds to random action choices (RC). The random initialization of the parameter optimization procedure introduces a source of noise, and the small error bars indicate the standard error of the mean over different runs of optimization (Methods, statistical model analysis). **B.** The expected posterior model probability [44, 45] given the whole dataset (Methods) with random effects assumption on the models. **C.** Accuracy rate of actions predicted by SurNoR (left scale and purple bars: mean and the standard error of the mean across participant) and the average uncertainty of SurNoR (right scale and dashed grey curve: entropy of action choice probabilities).

Given the behavioral data of all 12 participants, we estimated the log-evidence of all 13 algorithms, including SurNoR (see Methods). Comparison of the algorithms’ log-evidence (Fig. 4A) shows that SurNoR explains human behavior significantly better than its alternatives. In addition, a Bayesian model selection approach with random effects [44, 45] indicates that the SurNoR algorithm outperforms the alternatives with a protected exceedance probability of 0.99 (Fig. 4B and Methods).

The 1st episode of the 1st block is ideally suited to study how novelty influences behavior (middle panel in Fig. 4A). Our results show that all algorithms with novelty-seeking (+N) explain the behavior significantly better than models with random exploration strategy (RC) or optimistic initialization (MB+S+OI, MF+OI, and Hyb+S+OI), i.e., two classic approaches for exploration [17]. Our results also show that novelty-seeking explains behavior better than uncertainty-seeking (+U), a state-of-the-art exploration method in reinforcement learning [38, 39]. The models with uncertainty-seeking (MB+S+U, MF+S+U, and Hyb+S+U) use surprisal (i.e., the logarithm of our surprise measure) as an intrinsic reward as opposed to our model of novelty-seeking that uses novelty of states as an intrinsic reward.

As an alternative to novelty-seeking, participants might also solve the task simply by detecting and avoiding trap states. If so, the behavior of the participants can be explained if we replace the continuous novelty signal by a simple intrinsically generated binary signal equivalent to a negative reward. To address this issue, we tested two modified versions of the SurNoR algorithm (‘Binary Novelty’, see Supplementary Material). The 1st modification detects those states that have been encountered more often than some threshold value and assigns a fixed negative reward to them. The 2nd modification considers the *n* most frequently encountered states as bad states and, similar to the 1st modification, assigns a fixed negative reward to them – where *n* is a free parameter of the algorithm. Note that in both control algorithms, the constant negative rewards are treated as an intrinsic motivation signal – similar to novelty in SurNoR-algorithm except that the signal is a binary one. We estimated the log-evidence for both control algorithms. Our results show that SurNoR outperforms the 1st control algorithm by a 244 ± 11 difference in total log-evidence and by a 235 ± 5 difference in the log-evidence of the 1st episode of block 1, and outperforms the 2nd control algorithm by a 240 ± 11 difference in total log-evidence and by a 234 ± 5 difference in the log-evidence of the 1st episode of block 1. This observation rejects the hypothesis that participants simply identify ‘bad’ states by some binary signal.

Surprise becomes important in the 1st episode of block 2 (right panel in Fig. 4A). Indeed, the SurNoR model is significantly better than a hybrid model with novelty but without surprise (Hyb+N); similarly, model-free reinforcement learning with novelty and surprise (MF+S+N) is significantly better than model-free reinforcement learning with novelty alone (MF+N, right panel in Fig. 4A). Our results show that a constant adaptation rate as implemented in standard models without surprise is not sufficient to explain the choices of participants in the episode after the swap. Rather, the rate of learning and forgetting has to be modulated by a measure of surprise.

Overall, SurNoR is better than all 12 competing algorithms by a large margin, indicating that a combination of model-based and model-free algorithms explains behavior better than each algorithm separately, consistent with the notion of parallel, model-based and model-free, policy networks in the brain [3, 5, 6]. Going beyond these earlier studies, our results with SurNoR indicate that surprise and novelty are both necessary to explain human behavior in our task. Novelty is necessary to explain behavior during phases of exploration while surprise is necessary to explain behavior during the rapid re-adaption after a change in the environment.

### Individual decisions are dominated by the model-free policy network

Going beyond the relative merits of SurNoR in comparison with its alternative, we wondered whether the SurNoR model is also able to predict the individual actions of participants. Considering the most probable action of the model in a given state as the prediction of a participant’s next action in that state, we found that the SurNoR algorithm predicted the correct action in the 1st episode of the block 1 with an accuracy of 51±3% (3-fold cross validated, mean ± standard error of the mean over 12 participants, see Methods – Fig. 4C). Note that this accuracy is achieved in the absence of any a-priori preference of actions at initialization and is significantly higher than the accuracy rate of the naive random exploration strategy (25%, chance level); SurNoR’s predictions are also significantly better than the predictions of directed exploration through optimistic initialization (OI) of values or uncertainty-seeking policy (U) in a model-free, model-based, or a hybrid reinforcement learning algorithm without a novelty preference. Models with OI could at best predict 36±3% of the actions (for MB+S+OI), and the uncertainty-seeking strategy could at best predict 46±3% (for Hyb+S+U); one-sample t-test p-values for comparing their accuracy rates versus SurNoR’s are 0.0025 and 0.01, respectively. A crucial difference between OI and noveltybased exploration is that OI prefers those actions that have been less frequently chosen in the past, while novelty-seeking prefers actions that lead the agents to novel states, even if these are a few actions ahead and the outcome of the current action is known. Uncertainty-seeking is similar to OI because the uncertain actions are also those that have been less frequently chosen in the past.

Similarly, in the 1st episode of block 2, after the swap of states 3 and 7, the SurNoR algorithm predicts 56 ± 3% of the actions of the 12 participants (Fig. 4C). In the remaining episodes 2-5 of the two blocks, the SurNoR algorithm predicts 89 ± 2% of the action choices (Fig. 4C). Most of these actions move participants closer to the goal. The intrinsic uncertainty of action choices with the SurNoR model can be estimated from the entropy of the action choice probabilities across the four possible actions (Fig. 4C). Uncertainty decreases during the first three episodes as participants become familiar with the environment, but it jumps back to higher values after the swap of states at the beginning of block 2.

The SurNoR model combines a model-free with a model-based learning component (Fig. 5B), and we wanted to analyze the relative importance of each of the two components in explaining the action choices of participants. We therefore fitted the parameters of the SurNoR algorithm to the behavior of 12 participants (see Methods). In order to evaluate the relative importance of the two components, we normalized the Q-values of both branches and determined the relative weight of each branch (see Methods) during the 1st episode and 2nd-5th episodes of each block (Fig. 5D). We find that the model-free branch dominates the actions. Thus the world-model is of secondary importance for action selection and is mainly used to detect surprising events. In order to quantify the influence of surprise on learning, we plot the learning rate (of the model-free Q-values *Q_MF,R_* and *Q_MF,N_*) as a function of surprise (Fig. 5A). We find that non-surprising events lead to a small learning rate of 0.06 whereas highly surprising events induce a learning rate that is more than 8 times higher (Fig. 5A and Fig. 5C) indicating that surprise strongly influences the update of model-free Q-values.

**Figure 5:**
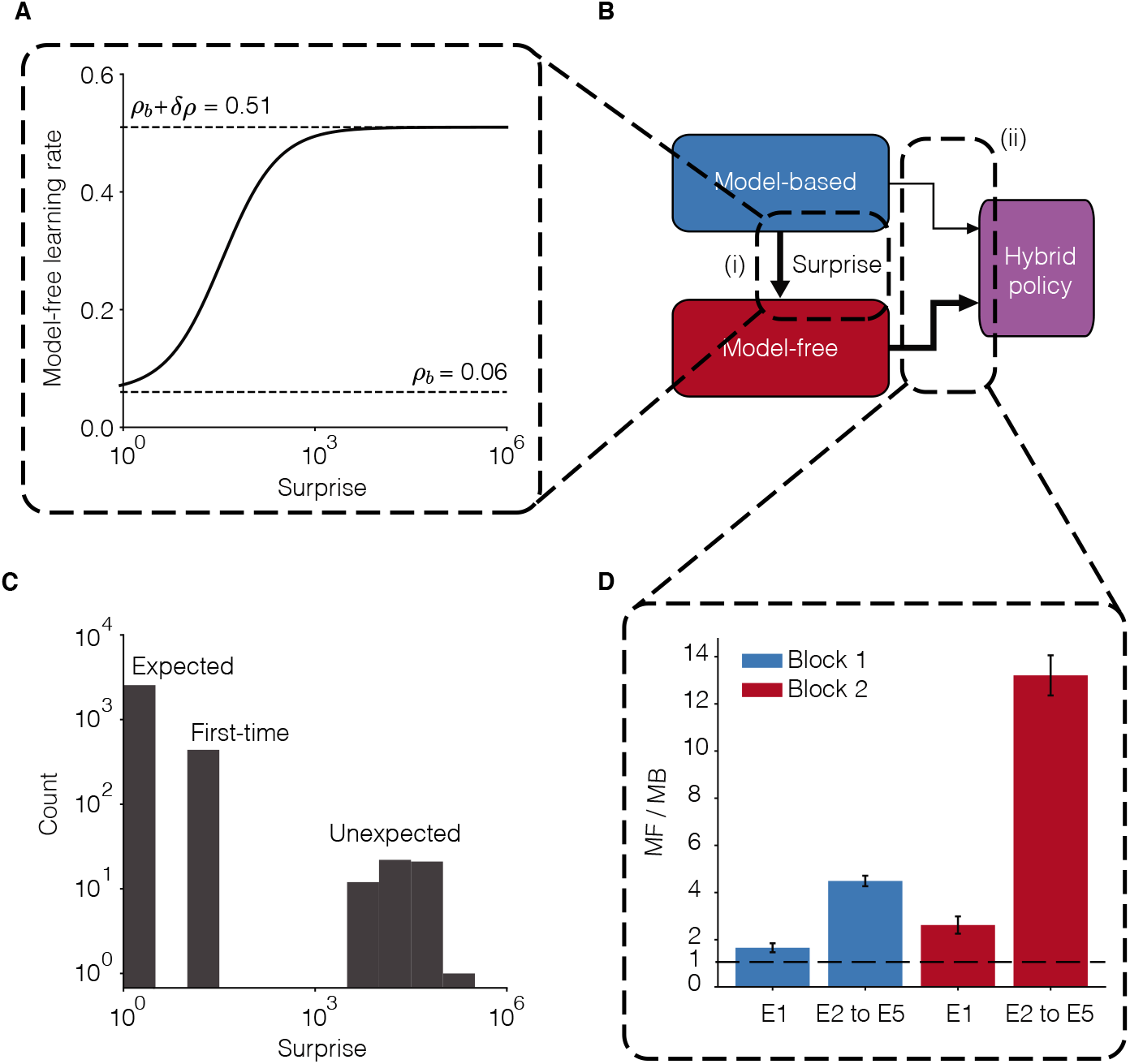
Model-based surprise modulates model-free learning. **A.** The learning rate of the model-free branch as a function of the model-based surprise, after fitting parameters to the behavior of all participants (see Eq. 9 in Methods). The model-free learning rate for highly surprising transitions is more than 8 times greater than the one for expected transitions. **B.** Three modules from the block diagram of Fig. 3C. There are two types of interactions between the model-based and the model-free branches of SurNoR: (i) The model-based branch modulates the learning rate of the model-free branch and (ii) the weighted (arrow thickness) outputs of the model-based and the model-free branches influence action selection (hybrid policy). **C.** The histogram of surprise values across all trials of 12 participants. The distribution is multimodal with high surprise for the unexpected transitions in the 1st episode of block 2, medium surprise for whenever a transition is experienced for the first time, and low surprise for the expected transitions. **D.** The relative importance of model-free (MF) compared to model-based (MB) in the weighting scheme of the hybrid policy during different episodes. Vertical axis: dominance of model-free (see Methods). Values larger than one (dashed line) indicate that the model-free branch dominates action selection. Error-bars indicate the standard error of the mean.

In conclusion, the SurNoR algorithm is able to predict individual actions with a high accuracy: it predicts 63 ± 2% of all actions and 74 ± 3% of the actions after the first time finding the goal. Our results suggests that participants (i) rely on propagation of novelty information via NPE in the first episode, (ii) base their decisions mainly on the model-free learner, and (iii) use surprise to modulate the learning rate.

### EEG correlates with novelty and surprise

Since surprise and novelty turned out to be important components of SurNoR in explaining the participants’ behavior, we wondered whether they are both reflected in the ERP. We first performed a grand correlation analysis in which we pooled the more than 2500 trials of 10 participants together after normalizing their ERPs to unit energy (see Methods; two participants were excluded because of noise artifacts in the recordings). We then computed the correlation of the ERP amplitudes, for each time point after the trial onset, with the model variables ‘Surprise’, ‘Novelty’, ‘Reward, ‘NPE’, or ‘RPE’ (capital initial letters indicate the 5 model variables). We find that Surprise, Reward, and RPE show significant positive correlations with the ERP amplitudes at around 300ms after stimulus onset (Fig. 6), in agreement with the well known correlation of the P300 amplitude with Surprise [21, 42, 46] and the well known correlation of the Feedback-Related Negativity (FRN) component with RPE [47, 48]. Moreover Novelty and NPE have, compared to Surprise, a broader positive correlation window with the ERP starting at around 200ms and ending at around 320ms after stimulus onset, and a second window with significant negative correlations from around 450ms to 550ms. Thus, Novelty and NPE have an ERP signature that is distinct from that of Surprise, Reward, or RPE (Fig. 6).

**Figure 6:**
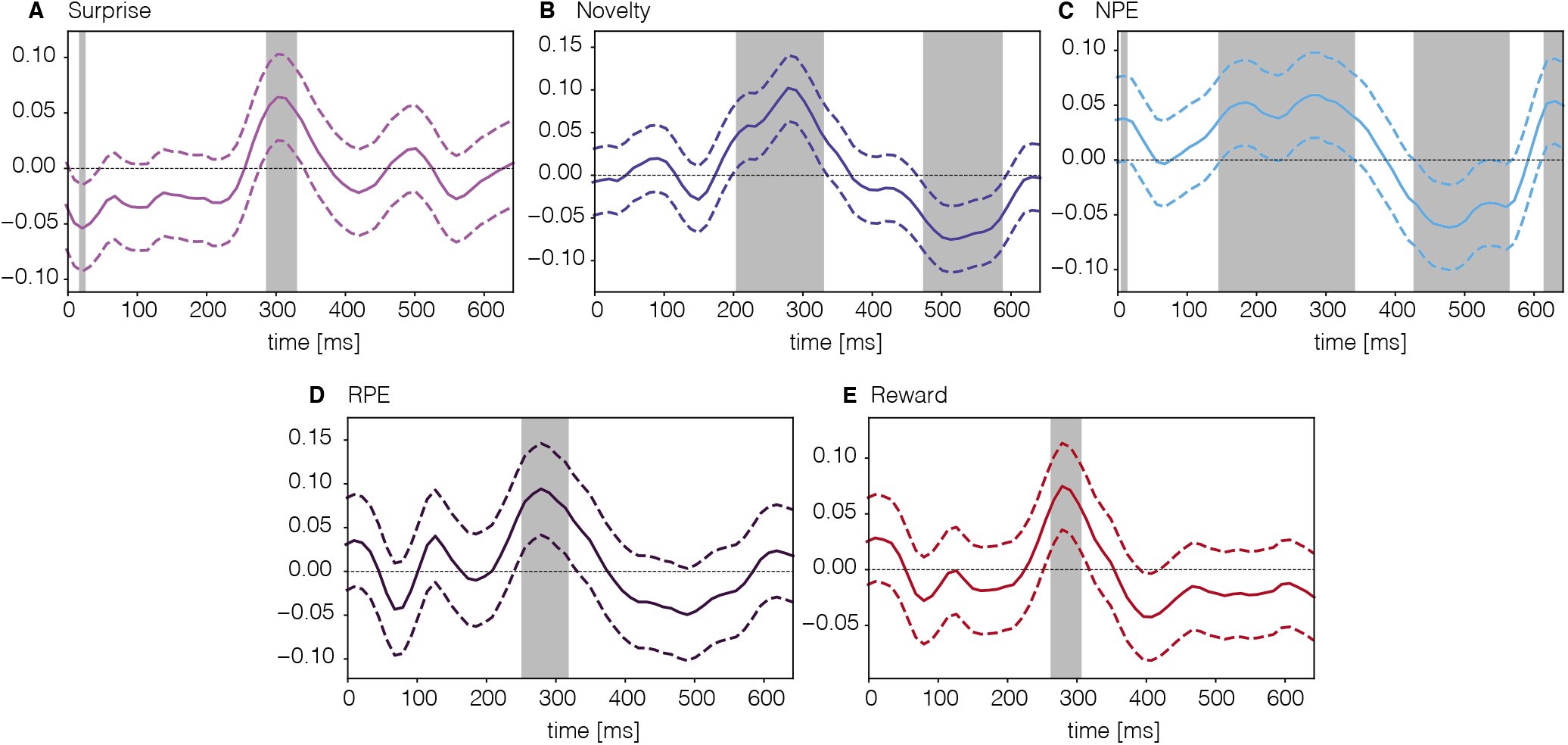
Grand correlation analysis of normalized ERPs over all 2524 trials of 10 participant. The dashed lines show confidence intervals. Shaded areas indicate intervals of significant correlations (FDR controlled by 0.1, one-sample t-test). Correlations of ERP with **A.** Surprise, **B.** Novelty, **C.** NPE, **D.** RPE (computed after excluding the trials from the 1st episode of the 1st block during which RPE is equal to 0) and **E.** Reward.

Second, we wondered how much of the variations in the ERP amplitudes could be explained by a linear combination of our five model variables, i.e., Suprise, Novelty, NPE, RPE, and Reward. We performed a trial-by-trial multivariate linear regression (MLR), separately for each participant. To be able to more precisely identify the separate contributions of each model variable to the regression, we needed to decorrelate them from each other. As expected from the design of the experiment, the cross-correlations between the normalized (zero mean and unit variance) sequences of Surprise, Novelty, and NPE are negligible (see Supplementary Material); however, the sequences of Reward and RPE are highly correlated with each other, mainly because Reward and RPE are both high at the goal state. Using principal components analysis over Reward and RPE, we find *R*_+_ (the sum of RPE and Reward) and *R*_ (their difference) as their decorrelated combinations (see Methods). We then extracted the components of Surprise, Novelty, and NPE orthogonal to *R*_+_ and *R*_ (see Methods). The resulting variables, denoted by an index⊥, are each orthogonal to *R*_+_ and *R*_, while staying very similar to the original signals, e.g., Surprise_⊥_ is highly correlated with Surprise, and NPE_⊥_ is highly correlated with NPE (see Supplementary Material).

For each participant, we considered the normalized Surprise_⊥_, Novelty_⊥_, NPE_⊥_, *R*_+_, and *R*_ as explanatory variables in order to predict the ERP amplitude at a given time point. We found 4 time intervals with an encoding power (adjusted R-squared, see Methods) significantly greater than zero (one-sample t-test, FDR controlled by 0.1, Fig. 7A and B; note that the adjusted R-squared can take negative values, e.g., see baseline in Fig. 7A). The 1st time window is around 193 ± 5ms; the P300 component can be linked to the 2nd time window which spans from 286 to 321 ± 5ms; since the 3rd time interval is long (from 392 to 487 ± 5ms), we split it into two time windows of equal size (W3a and W3b in Fig. 7B); and the last window extends from 532 to 574 ± 5ms.

**Figure 7:**
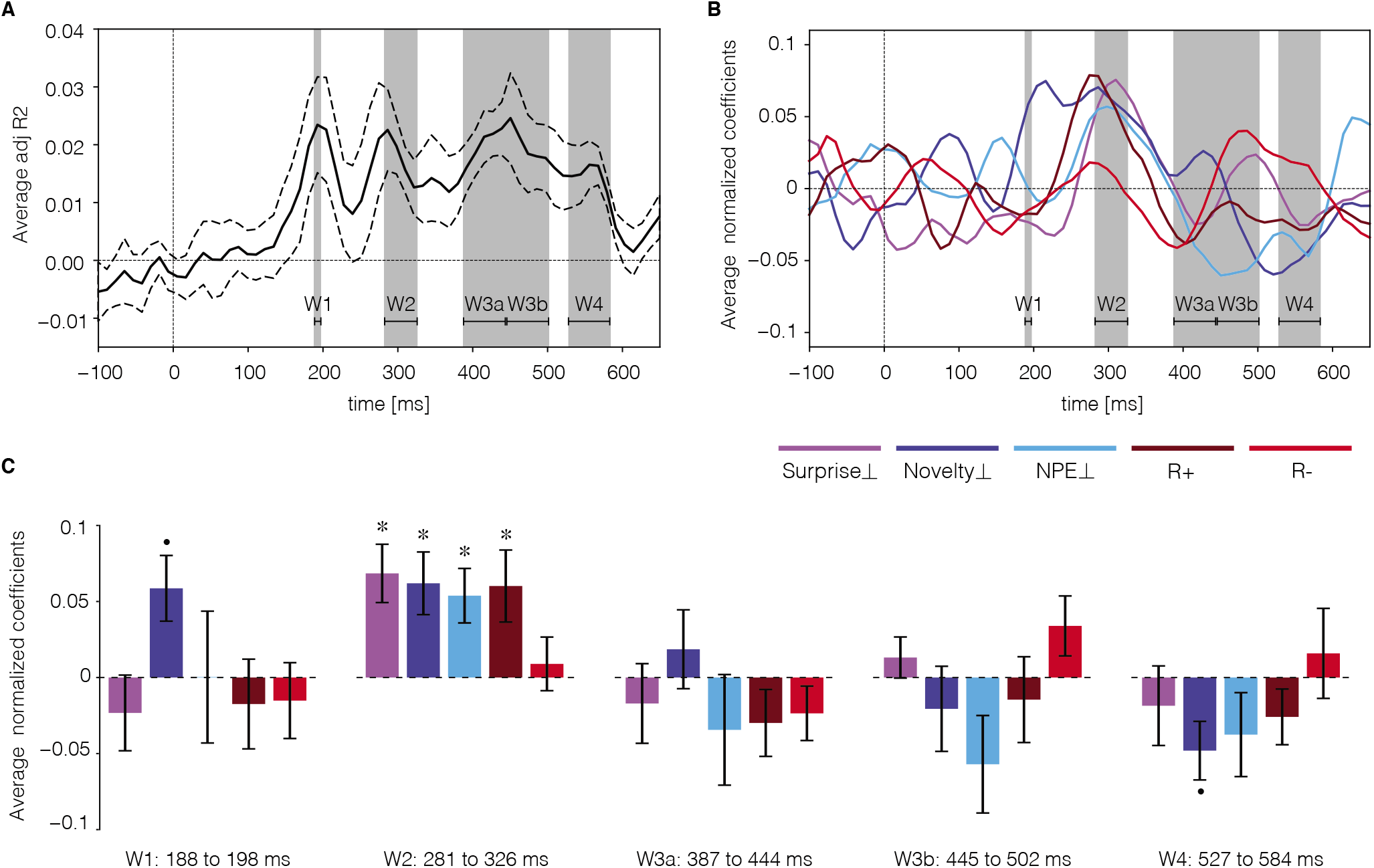
ERP variations explained by trial-by-trial and participant-by-participant multivariate linear regression analysis. Surprise_⊥_ (magenta), Novelty_⊥_ (dark blue), NEP_⊥_ (light blue), *R*_+_ x(brown) and *R*_–_ (red) were used as explanatory variables, and the ERP amplitude at each time point was considered as the response variable. **A.** Encoding power (adjusted R-squared values) averaged over 10 participants (dashed lines show the standard error of the mean) at each time point. Shaded areas and horizontal lines indicate four time intervals (W1,…, W4) of significant encoding power (FDR controlled by 0.1, one-sample t-test, only for the time-points after the baseline). The 3rd time interval has been split into two time windows of equal length for the analysis in C. **B.** Values of the regression coefficients (averaged over participants) for Surprise_⊥_, Novelty_⊥_, NEP_⊥_, *R*_+_, and *R*_–_ as a function of time. Errors are not shown to simplify the illustration. **C.** In each of the 5 time windows, the regression coefficients plotted in B have been averaged over time. Error bars show the standard error of the mean (across participants). Asterisks show significantly non-zero values (FDR controlled by 0.1 for each time window, one-sample t-test). The Novelty_⊥_ coefficients in the 1st and the last time windows (dot) have p-values of 0.03 and 0.04, respectively, which are not significant after FDR correction. In the second time window, Surprise_⊥_, Novelty_⊥_, NEP_⊥_, and *R*_+_ have significantly positive coefficients.

To study the contribution of surprise and novelty to encoding power, we focused on these time windows and tested the average regression coefficients of all explanatory variables in each time window in a second level analysis (Fig. 7C). Our results show that in the the second time window (286 to 321±5ms), Surprise_⊥_, Novelty_⊥_, NPE_⊥_, and *R*_+_ all have a significant positive regression coefficient in MLR (Fig. 7C, 2nd panel, FDR controlled by 0.1). While the coefficients for Surprise_⊥_, NPE_⊥_, and *R*_+_ sharply peak at around 300ms, the coefficient for Novelty_⊥_ has a broader peak starting at around 200ms (Fig. 7B) with a close to significant positive value during the 1st time window. This observation suggests that positive correlations of novelty with the ERP potentially extend from the 1st time window to the 2nd one, in agreement with our grand correlation analysis.

While consistent with previous studies of surprise in the ERP [20, 21, 42, 46], our results indicate that Surprise_⊥_ and Novelty_⊥_ contribute each separately to the ERP components at around 300ms. Furthermore, we find that NPE_⊥_ is yet another independent contributor to these components. As expected from previous studies [47, 48], *R*_+_ also shows a positive correlation with the ERP amplitude at around 300ms. While the multivariate analysis based on the 5 explanatory variables shows significance in the later time windows (Fig. 7A), individual contributions of Surprise_⊥_ or Novelty_⊥_ or NPE_⊥_ alone remain below significance level even though Novelty_⊥_ has a close to significant negative coefficient in the last window (Fig. 7C).

To summarize, the grand correlation analysis yields time windows of significance for Novelty and NPE that start 50 to 100ms *before* those of Surprise or Reward, indicating distinct contributions. Moreover, Novelty_⊥_ and NPE_⊥_ explain a significant fraction of the variations of the ERP at around 300ms that is not explained by Surprise_⊥_ and *R*_+_ alone. Importantly, NPE has significant correlations both in the grand correlation analysis and in the regression analysis, consistent with our earlier finding that NPE is important to explain behavior.

## Discussion

Combining a deep sequential decision-making task with the SurNoR model, an augmented reinforcement learning algorithm, we were able to extract the distinct contributions of surprise, novelty, and reward to human behavior. We found that the human brain (i) uses surprise to adapt their behavior to changing environments by modulating the learning rate and (ii) uses novelty as an intrinsic motivational drive to explore the world. Moreover, the model variables Suprise, Novelty, NPE, Reward, and RPE could well explain variations of the EEG amplitudes on a trial-by-trial basis.

As expected from previous theoretical [26–29, 31, 49] and experimental work [18, 22–24], our results suggest that the human brain uses surprise to modulate the learning of its world-model. Rather unexpectedly, however, our results indicate that humans hardly use their world-model to plan behavior; instead they mainly rely on model-free TD learning with eligibility traces to choose their next actions. Importantly, although the surprise signal is triggered by a mismatch between an observation and the predictions of the world-model, the modulatory effect of surprise is not limited to readjusting the world-model but also used to modulate the learning rate of model-free TD-learning. Following the common interpretations of model-based reinforcement learning algorithms as descriptions of human planning behavior and model-free reinforcement learning algorithms as descriptions of human habitual behavior [3, 5, 6, 50], our results suggest that (i) in the absence of surprise, humans prefer habitual behavior (potentially to reduce computational costs of decisionmaking [51, 52]) and (ii) errors in their world-model make them reconsider their habitual behavior. Our results extend findings that humans use hybrid policies in two-stage decision tasks [3, 5] to the case of deep sequential decision tasks in the presence of abrupt changes. The dominance of model-free behavior in such deep tasks does not exclude that humans used model-based planning in shallow tasks that are easily comprehensible thanks to a spatial arrangement of states or explicit instructions [53].

In the SurNoR model, surprise measures how *unexpected* the next event is, conditioned on the current state and the chosen action; in contrast, novelty measures how *rare* the next event is, independent of our expectations derived from a world-model. Our results show that exploration based on novelty-seeking (as opposed to random exploration [54], optimistic initialization [17], or uncertainty-seeking [38, 39]) can explain human behavior in our sequential decision-making task better than all tested alternative models. The presence of trap states in our behavioral paradigm makes novelty an ‘internally rewarding’ signal that helps participants to avoid ‘traps’. In general, an agent’s desire for seeking novel events depends on its inductive biases [9]. Following the idea of information search in active sampling [55, 56], we speculate that whether novelty is an informative cue (e.g., about the location of the goal) or not must be itself inferred by participants through the exploration procedure. Formulating and testing such a hypothesis is an interesting direction for future studies.

In contrast to published exploration strategies which give preference to those actions for which the outcome is most uncertain, i.e., those that have been tried least [38, 39, 54, 57], exploration based on novelty-seeking gives preference to actions that ultimately lead the agent to previously unvisited or less visited states, even if the agent is perfectly sure about the transition to the next state. Since, in the SurNoR model, Novelty is treated completely parallel to an external Reward, TD-learning based on NPE along with eligibility traces rapidly diffuses information about novel states to far-away non-novel states just as TD-learning based on RPE along with eligibility traces rapidly diffuses information about rewarding states to far-away non-rewarding states. Importantly, we found that NPE is a variable that has explanatory power for the ERP which adds to, and is different from, the explanatory power provided by Reward, RPE, or Surprise. The manifestation of a separate NPE signal in neural activities may open a new door for further developments of theories and experiments on novelty-driven activity of dopaminergic neurons and other neuromodulators [58–61].

The SurNoR algorithm suggests that participants treat novelty and reward as separately estimated values – as opposed to adding them into a single value estimator [10, 15, 16]. This separation enables participants to rapidly switch from exploration to exploitation, once they have found the goal. Based on this insight, we make the following prediction: if participants find a goal state but expect a second more rewarding goal state, they will continue to explore and potentially spend a large amount of time in a novelty-rich segment of an extended version of the environment of Fig. 1 (see Supplementary Material).

Our EEG analysis shows that variables of the SurNoR model can significantly explain the variations of ERP amplitudes in several time-windows: Surprise, Novelty, NPE, and Reward/RPE all significantly contribute to the encoding power in the time-window around 300ms. The positive contributions of Novelty, Surprise, and Reward/RPE in this time-window are consistent with previous studies of the P300 and the FRN component [20, 21, 35, 42, 46–48]. Whereas in earlier studies contributions of Novelty, Surprise, and Reward were often mixed together [20, 21, 35, 42, 46–48], we have shown here separate, additive contributions of these three variables as well as a further contribution of NPE. The effect of Novelty appears in ERPs earlier (at around 200ms) than the correlations with the other variables; moreover, contributions of Novelty are distinct from those of Surprise in the time window after 400ms.

In conclusion, surprise and novelty are conceptually distinct concepts that also give rise to different temporal components in the ERP. Our results suggest that humans use novelty-seeking for efficient exploration and surprise for a rapid update of both their internal world-model and their model-free habitual responses.

## Methods

### Experimental setup

Stimuli were presented on an LCD screen that was controlled by a Windows 7 PC. Experiments were scripted in MATLAB using the Psychophysics Toolbox [62].

### Participants

14 paid participants joined the experiment. Two participants quit the experiment, hence, we analysed data for 12 participants (5 females, aged 20-26 years, mean = 22.8, sd = 1.7). All participants were right-handed and naive to the purpose of the experiment. All participants had normal or corrected-to-normal visual acuity. All participants provided written consent. The experiment was approved by the local ethics committee.

### Stimuli and general procedure

Before starting the experiment, we showed the participants the goal image that they were required to find on a computer screen. Next, participants were presented, in random order, all the other images that they might encounter during the experiment. Thereafter, participants clicked the ‘start’ button to start the experiment. At each trial, participants were presented an image (state) and four grey disks below the image. Clicking on one of the disks (action) led participants to a subsequent image; for details of timing see Fig. 1A. Participants clicked through the environment until they found the goal state which finished the episode. An episode *n* started at a random state *i*(*n*) which was the same for all participants; in our experiment we used *i*(1) = 6, *i*(2) = 9, *i*(3) = 4, *i*(4) = 5, and *i*(5) = 8.

### EEG recording and processing

EEG signals were recorded using an ActiveTwo Mk2 system (BioSemi B.V., The Netherlands) with 128 electrodes at a 2048Hz sampling rate. Two participants were excluded from EEG analysis because of their noisy and low quality signals caused by substantial movements during the experiment. Data were band pass filtered from 0.1Hz to 40Hz and down sampled to 256Hz. EEG data were recorded with a Common Mode Sensor (CMS) and re-referenced using the common average referencing method. We used EEGLAB [63] toolbox in MATLAB to perform the EEG preprocessing. We extracted EEG trials from 200ms before to 700ms after the state onset. Trials in which the change in voltage at any channel exceeded 35 *μ*V per sampling point were discarded. Eye movements and electromyography (EMG) artefacts were removed by using independent component analysis (ICA). The baseline activity was removed by subtracting the mean calculated over the interval from 200ms to 0ms before the state onset. EEG data of selected prefrontal electrodes (Fz, F1, F2, AFz, FCz) were averaged for ERP analysis. We further smoothed (moving averaging with the window of length 50ms) and downsampled (to the sampling rate of 1 sample per 11.7ms) ERPs. Data were analyzed during the time window from 0 to 650ms after state onset (blue interval in Fig. 1A). For multivariate regression analysis, a 100ms-baseline was also included for sanity check. As a result, each trial (from 100ms before to 650ms after the onset of the state) consisted of 65 time points.

### SurNoR algorithm

We present a more detailed formulation and the psudocode of the SurNoR algorithm in Supplementary Material. Here we outline the algorithm in brief.

We formally define the **Novelty** of a state *s* at time *t* as 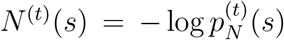, where 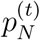 is defined in Eq. 2; see Supplementary Material for further discussion. When observing the image corresponding to state *s*_*t*+1_ at time *t* + 1, after taking action *a_t_*, the novelty *n*_*t*+1_ = *N*^(*t*)^(*s*_*t*+1_) is treated as an internal novelty-reward, completely analogous to the treatment of external rewards in reinforcement learning. This analogy between external reward and novelty is inspired by earlier experimental studies [58, 59, 64]. As a result, at time *t* + 1, agents receive three signals: the next state *s*_*t*+1_, the external reward *r*_*t*+1_ (i.e., the indicator of whether *s*_*t*+1_ is the goal state or not), and the novelty *n*_*t*+1_ (indicated as the output of the grey block in Fig. 3A).

The SurNoR algorithm has two branches, i.e., a model-based and a model-free one, which interact with each other (Fig. 3A, blue and red blocks). The **model-based branch** computes the Bayes Factor Surprise [27]

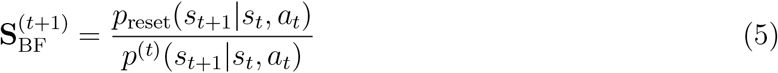

where *p*^(*t*)^(*s*_*t*+1_|*s_t_*, *a_t_*) is the probability of observing *s*_*t*+1_ by taking action *a_t_* in state *s_t_* as estimated from the current world-model (cf., Eq. 4), and *p*_reset_(*s*_*t*+1_|*s*_*t*_, *a_t_*) is the probability of observing *s*_*t*+1_ by taking action *a_t_* in state *s_t_* with the assumption that the environment has experienced an abrupt change between time *t* and *t* +1, so that the world-model should be reset to its prior estimate. In this work, we assume that the prior estimate *p*_reset_(*s*_*t*+1_ |*s_t_*, *a_t_*) = 1/11 is a uniform distribution over states and hence constant as stated in Eq. 3; see Supplementary Material and [27] for further discussion. Note that in Fig. 3A, Fig. 3B, and Fig. 5 we suppressed the factor 1/11 and directly plotted 1*/p*^(*t*)^(*s*_*t*+1_|*s_t_*,*a_t_*) as the surprise value. As an aside we note that since the state prediction error [5] is defined as *SPE*_*t*+1_ = 1 – *p*^(*t*)^(*s*_*t*+1_|*s_t_*, *a_t_*), the Bayes Factor Surprise can be written as 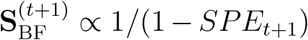. The definition of the Bayes Factor Surprise is valid for arbitrary volatile environments [27]. However, since in our experimental setting *p*_rese_,(*s*_*t*+1_|*s_t_,a_t_*) is assumed to be uniform, the Bayes Factor Surprise **S**_BF_ is a monotone function of Shannon Surprise and hence comparable to previous studies [21, 42, 43].

The value 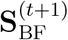 is used in the model-based branch to update the world-model using the Variational SMiLe algorithm [27], an approximate Bayesian learning rule with surprise-modulated learning rate designed for volatile environments with abrupt changes. Updating the world-model is equivalent to updating the pseudo-counts 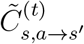, introduced in Eq. 4, for all possible *s, a*, and *s*’. The Variational SMiLe algorithm [27] yields the updates

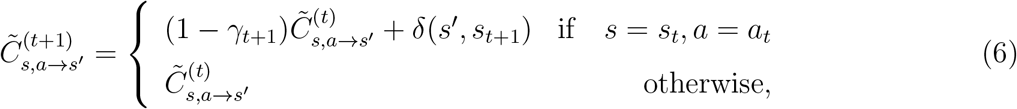

where *δ* is the Kronecker delta function, and *γ*_*t*+1_ is the surprise modulated adaptation factor [27]

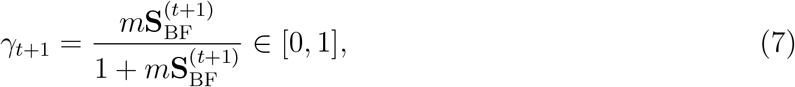

with *m* ≥ 0 a free parameter related to the volatility of the environment [27]. Note that if the transition from *s* to *s*’ caused by action *a* is unsurprising, then the pseudo-count of that transition is increased by one (because *γ* = 0 for **S**_BF_ = 0). However, if this transition has a high surprise, the earlier pseudo-count is reset to zero (because *γ* → 1 for **S**_BF_ → ∞) and the observed transition is counted as the first one. The updated world-model is then used to update a pair of *Q*-values, i.e., 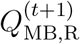. for Reward and 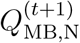 for Novelty, by solving the corresponding Bellman equations with a variant of prioritized sweeping [17, 65, 66]; see Supplementary Material for details.

The **model-free branch** computes Reward and Novelty prediction errors, *RPE*_*t*+1_ and *NPE*_*t*+1_. As usual, RPE is defined as *RPE*_*t*+1_ = *r*_*t*+1_ + *λ_R_V*_MF,R_(*s*_*t*+1_) – *Q*_MF,R_(*s_t_,a_t_*), where *λ_R_* is the discount factor for reward, and *V*_MF,R_(*s*_*t*+1_) = max_*a*_ *Q*_MF,R_(*s*_*t*+1_,*a*) is the value of the state *s*_*t*+1_. Analogously, NPE is defined as *NPE*_*t*+1_ = *n*_*t*+1_ + *λ*_N_*V*_MF,n_(*s*_*t*+1_) – *Q*_MF,N_(*s_t_,a_t_*), where *λ_N_* is the discount factor for novelty, and *V*_MF,N_(*s*_*t*+1_) = max_*a*_ *Q*_MF,N_(*s*_*t*+1_, *α*) is the novelty value of the state *s*_*t*+1_.

A Surprise-modulated TD-learner with eligibility traces is used for updating the two separate sets of *Q*-values. To have the most general setting, two separate eligibility traces are used for the update of *Q*-values, one for reward 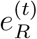 and one for novelty 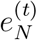. The eligibility traces are initialized at zero at the beginning of each episode. The update rules for the eligibility traces after taking action *α_t_* at state *s_t_* is

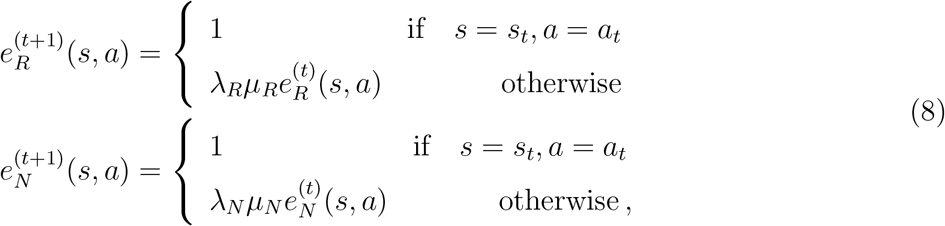

where *λ_R_* and *λ_N_* are the discount factors defined above, and *μ_N_* ∈ [0,1] and *μ_R_* ∈ [0,1] are the decay factors of the eligibility traces for novelty and reward, respectively. The update rule is then 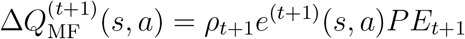, where *e*^(*t*+1)^ is the eligibility trace (i.e., either 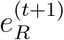 or 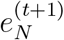)), *PE*_*t*+1_ is the prediction error (i.e., either *RPE*_*t*+1_ or *NPE*_*t*+1_) and

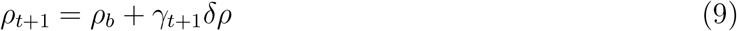

is the surprise-modulated learning rate with parameters *ρ_b_* for the baseline learning rate and *δ_ρ_* for the effect of Surprise.

Finally, actions are chosen by a hybrid policy (Supplementary Material) using a softmax function of a linear combination of the values 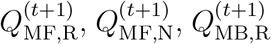, and 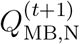 (the purple block in Fig. 3.A), similar, but not identical to [5, 6]. The weight of 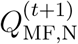 and 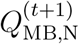 is non-zero only in the 1st episodes of blocks 1 and 2. Overall, the SurNoR algorithm has 18 free parameters.

### Statistical model analysis and fit to behavior

In addition to SurNoR, we considered 12 alternative algorithms with 0 to 18 free parameters and a control algorithm for SurNoR with Binary Novelty with 19 free parameters (Supplementary Material). For each algorithm, we used 3-fold cross-validation and computed its maximum loglikelihood for each participant, similar to existing methods [8]: (i) we divided participants into 3 folds each consisting of four participants; (ii) for participant *i*, we estimated the parameters of the algorithm by maximizing the likelihood function of the folds which did not include participant *i;* and (iii) we computed the log-likelihood for participant *i* using the estimated parameters. The maximization procedure was done by coordinate ascent (using grid search for each coordinate); we repeated the procedure until convergence starting from 25 different random initial points. We further repeated the whole process 4 times to have an estimation of the variability resulting from random initialization of the optimization procedure. The error bars in Fig. 4A are calculated using these 4 samples.

Similar to studies in economics and statistics [67, 68], we considered, for each participant and each algorithm, the cross-validated maximum log-likelihood (averaged over the 4 repetitions) as the log-evidence [69]. The sum (over participants) of the log-evidences for each algorithm is shown in Fig. 4A – see [70] for a tutorial on the topic. As a convention, differences greater than 3 or 10 are considered as significant or strongly significant, respectively [69, 70]. The model posterior and protected exceedance probabilities in Fig. 4B are computed by using the participant-wise log-evidences (averaged over the 4 repetitions) and following the Bayesian model selection method of [44, 45] (available in SPM12 toolbox for MATLAB). We used a Dirichlet distribution with parameters equal to 1 over the number of models (1/13) as the prior distribution. This choice of prior is equivalent to stating that the prior information is worth as much as the observation coming from a single participant [69]; it is also a default choice of prior in the VBA toolbox [71].

The accuracy rate and the uncertainty in Fig. 4C are computed by the same cross-validation procedure. We define accuracy as the ratio of the number of trials with correctly predicted actions to the total number of trials; for a given trial, whenever the action taken by the participant had the maximum probability under the policy but shared with other *n* – 1 (e.g., 2) actions, we counted that trial as 1/*n* (e.g., 0.333) correctly predicted. With this procedure, the accuracy rate of the random choice algorithm is 25%. We define the uncertainty of one participant in an episode as the average of the entropy of his or her policy over all trials of that episode. Both the accuracy rate and the uncertainty were computed for each participant separately, but only the mean and the standard error across participants are reported in Fig. 4C.

For EEG analysis, we only considered the SurNoR algorithm (i.e., the winner of statistical model selection). To have the same set of parameters for all participants, we fitted our model to the whole behavioral data set (overall 3047 actions) by maximizing total log-likelihood – similar to [6]. For each of 100 random initialization points, maximization was implemented as coordinate ascent until convergence (using grid search for each coordinate). Amongst the 5 local maxima with high but not significantly (< 3) different log-evidence, we kept the model which had the greatest encoding power in multivariate regression analysis of EEG. The fitted parameters are reported in in Supplementary Material.

The plots in Fig. 5 corresponds to this set of parameters. Since the softmax operator of the hybrid policy has a free scale parameter, the effective weight of each branch of the hybrid policy in Fig. 5A (i.e., model-free and model-based) is computed as the fitted weight of each component times its average difference in Q-values. For example, 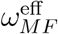 is equal to 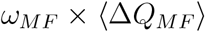, where *ω_MF_* is the weight of model-free Q-values in the hybrid policy and 〈Δ*Q_MF_*〉 is the average (over trials) of the difference between *Q_MF_* of the best and the worst actions. The weight 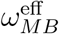 for the model-based branch is defined analogously. The dominance of the model-free branch is defined as 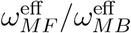.

### EEG Analysis

#### Participant-based regression analysis

Given *N* trials (across all episodes of both blocks) of a given participant, the matrix *X*_raw_ for this participant is an *N* by 5 matrix whose rows correspond to trials and whose columns correspond to normalized model variables (i.e., Surprise, Novelty, NPE, RPE, and Reward). For example, if the sequence of reward prediction error values for this participant is *z*_1:*N*_, then one column of the matrix *X*_raw_ is equal to (*z*_1:*N*_ – *μ_z_*)/*σ_z_* where *μ_z_* is the mean and *σ_z_* is the standard deviation of *z*_1:*N*_, and one row of the matrix *X*_raw_ is equal to the normalized values of Surprise, Novelty, NPE, RPE, and Reward for one trial. We constructed the feature matrix *X* from *X*_raw_ by applying the following steps: (i) we put 2 columns of *X* to be equal to normalized Reward plus RPE and Reward minus RPE, calling them R_+_ and R_–_, respectively; since Reward and RPE were normalized, their sum and difference correspond to their principal components (Supplementary Material); (ii) we orthogonalized each of the other variables to R_+_ and R_–_. For example, NPE_⊥_ is NPE minus its projection on R_+_ and R_–_, followed by a renormalization step (see Supplementary Material).

For each trial, time of the ERP is measured with respect to the image onset. For a given time point, we defined the target vector *y* as an *N* dimensional vector whose elements are equal to the normalized (zero mean, unit variance) amplitude of ERPs at that particular time point in different trials. Since we have 65 time points, the response matrix *Y* is a *N* by 65 matrix. We then performed multivariate linear regression (MLR) by considering *ŷ* = *Xβ* as an estimation of *y* and found *β* by ordinary least squared error minimization. The encoding power for a single time point and for the given participant was calculated as adjusted R-squared [72]. Note that adjusted R-squared can in principle be negative – which is the case for our regression analysis over baseline in Fig. 7A.

Fig. 7A shows the mean and the standard error of the mean of the encoding power over participants and for each time-point. The threshold for rejecting the null hypothesis is computed using the Benjamini and Hochberg algorithm [69] for controlling false discovery rate (FDR) by 0.1. Fig. 7B shows the average (over participants) of *β* values as a function of time. For Fig. 7C, we first average the *β* values over time within each time window, and then evaluate their mean and their standard error of the mean (over participant). The FDR correction was done separately for each time window.

#### Grand correlation analysis

Similar to the approach of [73], we pooled all trials of all participants together, i.e., we concatenated *X*_raw_s and *Y*s for different participants. However, before concatenation, to remove the difference in the between-participant variations of ERPs energy (i.e., 2nd moment), we divided ERPs of each participant by the overall squared-energy of that participant’s ERPs, i.e., we replaced *Y* by 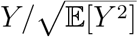. The correlations in Fig. 6 are computed between columns of concatenated *X*_raw_s and concatenated *Y*s. For RPE, we removed the trials corresponding to 1st episodes of the 1st blocks because RPEs are exactly equal to zero.

## Acknowledgement

AM thanks Johanni Brea and Vasiliki Liakoni for useful discussions on behavioral modeling and data analysis. This research was supported by Swiss National Science Foundation No. CRSII2 147636 (Sinergia, MHH and WG) and No. 200020 184615 (WG), and by the European Union Horizon 2020 Framework Program under grant agreement No. 785907 (Human Brain Project, SGA2, MHH and WG).

## Author Contributions

MHH, HAX and WG designed the experiment. HAX performed the experiments. AM designed the model and performed the data analysis. HAX and MPL developed preliminary models and data analysis. AM, HAX, MHH and WG wrote the paper.

