## Supplementary Material for "Novelty is not Surprise: Human exploratory and adaptive behavior in sequential decision-making"

Just as in the main text, RPE denotes the reward prediction error, NPE the novelty prediction error, TD-learning stands for Temporal Difference learning, and SurNoR stands for Surprise-Novelty-Reward. This supplementary material is structured as follows.

The details of the SurNoR model are given in Section 1 with pseudocode summarized in Algorihm 1-4. Section 2 and Table 1 present the alternative algorithms used in the main text.

In Section 3 we present the proof of the claim that $R_+ =$ Reward+RPE and $R_- =$ Reward−RPE are the principal components of normalized Reward and RPE. The correlation matrices between model-variables (i.e., Surprise, Novelty, NPE, Reward, and RPE) and the schematic of deccorrelating them is shown in Fig. S1 of Section 4.

The expected length of the 1st episode of block 1 under the random choice policy is calculated in Section 5. Section 6 formalizes our prediction made in the Discussion section of the main text. Table 2 in Section 7 lists the parameters of the SurNoR model after fitting to behavior and used for EEG analysis.

### 1 Surprise-Novelty-Reward (SurNoR) algorithm

The SurNoR algorithm (Alg. 1) combines surprise signals with novelty and reward so as to explore and learn the environment and exploit rewards. A simple block diagram of the algorithm is shown in Fig. 3C of the main text. In the SurNoR algorithm, a model-based and a model-free branch interact with each other. The output of each branch is a pair of $Q$-values for estimated novelty and estimated reward. The model-based branch updates model-based $Q$-values using a world-model that is estimated online, while the model-free branch uses a surprise-modulated TD-learner for updating the model-free $Q$-values. Finally, actions are selected following a hybrid policy that combines model-free and model-based $Q$-values - see [1, 2] for similar approaches. In this section, we describe the SurNoR algorithm in detail. For the sake of clarity and coherence, we repeat here some details already explained in the main text.

**Formalization of the environment.** The state and the action at time $t$ are random variables $S_t$ and $A_t$ which take values in the finite sets $\mathcal{S}$ and $\mathcal{A}$, respectively. In the particular case of our experiment, we have $\mathcal{S} = \{1, ..., 10, G\}$ and $\mathcal{A} = \{1, ..., 4\}$. Taking a Bayesian perspective, we consider the transition probability matrix as another random variable $\Theta$, i.e.

$$\mathbb{P}(S_{t+1} = s' | S_t = s, A_t = a, \Theta = \theta) = \theta_{s,a}(s'). \tag{S1}$$

where the values of $\theta_{s,a}(s')$ for combinations of $s$, $a$, and $s'$ are unknown and needed to be estimated from experience. Since our environment is deterministic, except for the switch of two states before the start of block 2, the real transition probabilities are

$$\theta^{\text{real}}_{s,a}(s') = \delta(s', T(s, a)), \tag{S2}$$

where $T(s, a)$ denotes the target state of the transition from state $s$ given action $a$, and the Kronecker $\delta$ is defined as $\delta(x, x') = 1$ if $x = x'$ and zero otherwise; $T(s, a)$ corresponds to the arrows in Fig. 1B and Fig. 1D in the main text. The target state depends on the block number in our experiment. Note that $T(s, a)$ is unknown to the participants and to the SurNoR algorithm.

**Definition of novelty.** While a participant moves in the environment, the count

$$C_s^{(t)} = |\{t' : 1 \leq t' \leq t \text{ and } s_{t'} = s\}|$$

indicates how often state $s$ has been encountered up to time $t$. We assume that at each time $t$, participants are able to estimate the empirical frequency $p_N^{(t)}(s)$ of encountering state $s \in \mathcal{S}$, formally defined as

$$p_N^{(t)}(s) = \frac{C_s^{(t)} + 1}{t + |\mathcal{S}|}, \tag{S3}$$

where $|\mathcal{S}|$ is the total number of states (i.e., 11 for our experiment). Note that the participants know the total number of states, due to the pre-experiment introduction. The empirical frequency in Eq. S3 is equal to the expected probability of observing state $s$ given $s_{1:t}$ under the assumption of a uniform prior over the probabilities of observing different states: before the start of the experiment all $|\mathcal{S}|$ states have the same prior probability $p_N^{(0)}(s) = 1/|\mathcal{S}|$.

We define the novelty of the state $s$ at time $t$ as the negative logarithm of the empirical frequency

$$N^{(t)}(s) = -\log(p_N^{(t)}(s)). \tag{S4}$$

In our algorithm, novelty acts just like an internally generated reward or exploration bonus (see subsection 'Formalizing model-based $Q$-values'). The main difference between our definition of novelty and most of the previously proposed measures of 'exploration bonus' [3–9] is in their dependency upon states *and* actions: while the usual exploration bonus measures are functions of state-action pairs, we define our novelty as a function of states only. Our choice is more consistent with the behavior of participants in our experiment, since a reasonable strategy of participants is to visit the states that they rarely encounter in the experiment as opposed to testing all actions in all states. From this perspective, our novelty measure is similar to the exploration bonus proposed by Bellemare, et al. (2016) [10].

In three of our alternative algorithms (see Section 2) we use a state-of-the-art exploration strategy [8, 9] which defines exploration bonus (or internal reward) as a function of the pairs of states and actions. We compare these algorithms with SurNoR (see Fig. 4 in the main text).

#### 1.1 Model-based branch of SurNoR

The pseudocode for the model-based branch is shown in Alg. 2. In this subsection, the details are explained.

**World-model.** The participants knew that there were 11 states and 4 possible actions in each state. However, they were not aware of the actual transition probability matrix. In particular, they did not know whether the environment is deterministic or stochastic. Therefore, we define a participant's model of the world as an approximation $q$ of the posterior distribution of the transition probability matrix, similar to the approach of [11–13],

$$q^{(t)}(\theta) \approx \mathbb{P}(\Theta = \theta | S_{1:t} = s_{1:t}, A_{1:t-1} = a_{1:t-1}). \tag{S5}$$

In the following, we call $q$ the belief of the participant. We assume that a participant estimates the transition probabilities by a weighted average

$$\hat{\theta}^{(t)} = \mathbb{E}_{q^{(t)}}[\Theta] = \int \theta q^{(t)}(\theta) d\theta, \tag{S6}$$

where the weighting factor is given by the belief $q^{(t)}$. For convenience, the transition probability $\hat{\theta}^{(t)}_{s,a}(s')$ is written as $p^{(t)}(s'|s,a)$ in the main text, e.g., in Eq. 3 and Eq. 4.

For exact Bayesian inference one needs to explicitly specify the generative model which governs the transition. Particularly, the dynamics of $\Theta$ over time should be known, e.g., whether it is fixed, continuously drifting, or experiencing abrupt changes [14–17]. However, rather than making explicit assumptions about the generative model as a starting point for exact Bayesian inference, we work with a general (parametric, see the next part) distribution $q^{(t)}$ which is updated by an appropriate learning algorithm after each observation, similar to approaches in machine learning [18, 19].

**Beliefs as Dirichlet distributions.** We assume that the transition probabilities from different state-action pairs are independent of each other, i.e.

$$q^{(t)}(\theta) = \prod_{s \in \mathcal{S}, a \in \mathcal{A}} q^{(t)}(\theta_{s,a}), \tag{S7}$$

where $\theta_{s,a}$ is defined as in Eq. S1. As a natural[1] choice for a probability distribution over transition probabilities, we consider the belief $q^{(t)}(\theta_{s,a})$ to be a Dirichlet distribution with parameter $\alpha^{(t)}_{s,a}$:

$$q^{(t)}(\theta_{s,a}) = \mathrm{Dir}(\theta_{s,a}; \alpha^{(t)}_{s,a}). \tag{S8}$$

As a result, at each time $t$, the belief of participants about their environment can be summarized in the set $\alpha^{(t)} = \{\alpha^{(t)}_{s,a}, \forall (s,a) \in \mathcal{S} \times \mathcal{A}\}$. We consider the parameter of the prior belief $q^{(1)}$ (i.e., $\alpha^{(1)}$) to be the same for all transitions, i.e.,

$$\alpha^{(1)} = \{\alpha^{(1)}_{s,a}(s') = \epsilon, \quad \forall (s, s', a) \in \mathcal{S} \times \mathcal{S} \times \mathcal{A}\}, \tag{S9}$$

where $\epsilon > 0$ is a free parameter. With this choice of prior, $\hat{\theta}^{(1)}_{s,a}$ (i.e., the prior estimate of the transition probabilities from the pair of state $s$ and action $a$) is a uniform distribution over states. Furthermore, the free parameter $\epsilon$ expresses how deterministic the transitions are from the point of view of a participant, i.e., smaller values of $\epsilon$ indicate a more deterministic interpretation of the environment.

Using a Dirichlet distribution for the belief $q^{(t)}$ and Eq. S6, a participant's estimation of the transition probabilities is

$$\hat{\theta}^{(t)}_{s,a}(s') = \frac{\alpha^{(t)}_{s,a}(s')}{\sum_{\tilde{s}' \in \mathcal{S}} \alpha^{(t)}_{s,a}(\tilde{s}')}. \tag{S10}$$

Note that, the pseudo-counts $\tilde{C}^{(t)}_{s,a \to s'}$ in Eq. 4 of the main text is equal to $\alpha^{(t)}_{s,a}(s') - \epsilon$.

**Definitions of surprise.** We work with the 'Bayes Factor' surprise $\mathbf{S}_{\mathrm{BF}}$ [14]. Consider the transition initiated at time $t$, i.e., $(S_t = s, A_t = a) \to (S_{t+1} = s')$ . The Bayes Factor surprise corresponding to this transition is [14]

$$\mathbf{S}^{(t+1)}_{\mathrm{BF}} = \frac{\hat{\theta}^{(1)}_{s,a}(s')}{\hat{\theta}^{(t)}_{s,a}(s')}. \tag{S11}$$

[1] If transition probabilities are stationary and have a uniform (or in general any Dirichlet) prior, exact Bayesian inference yields a Dirichlet distribution.

Due to the particular form of the prior $q^{(1)}$ that we chose, $\hat{\theta}^{(1)}_{s,a}(s')$ is constant. As a result, the surprise $\mathbf{S}^{(t+1)}_{\text{BF}}$ at time $t+1$ is proportional to the inverse of the estimated probability $\hat{\theta}^{(t)}_{s,a}(s')$ of the transition initiated at time $t$. In Eq. 5 of the main text, $\hat{\theta}^{(t)}_{s,a}(s')$ is written as $p^{(t)}(s'|s,a)$, and $\hat{\theta}^{(1)}_{s,a}(s')$ is written as $p_{\text{reset}}(s'|s,a)$.

We note that in the particular case of our behavioral paradigm, the Shannon surprise [20] is just the shifted logarithm of the 'Bayes Factor' surprise, i.e., $\mathbf{S}^{(t+1)}_{\text{Sh}} = \log \mathbf{S}^{(t+1)}_{\text{BF}} + \log|\mathcal{S}|$. Furthermore, the state prediction error (SPE) [1] is an increasing function of the 'Bayes Factor' surprise, i.e., $\text{SPE}^{(t+1)} = 1 - \frac{1}{|\mathcal{S}|\,\mathbf{S}^{(t+1)}_{\text{BF}}}$. Hence, surprise-modulated learning rates in the SurNoR algorithm can alternatively be expressed in terms of $\mathbf{S}^{(t+1)}_{\text{BF}}$ or $\mathbf{S}^{(t+1)}_{\text{Sh}}$ or $\text{SPE}^{(t+1)}$.

**Surprise-modulated update of the belief.** Learning the world-model corresponds to updating the parameters of the Dirichlet distribution after each transition. Consider the transition $(S_t = s, A_t = a) \to (S_{t+1} = s')$ initiated at time $t$ which generates a surprise $\mathbf{S}^{(t+1)}_{\text{BF}}$ at time $t+1$. The surprise-modulated adaptation rate is defined as [14]

$$\gamma(\mathbf{S}^{(t+1)}_{\text{BF}}, m) = \frac{m\mathbf{S}^{(t+1)}_{\text{BF}}}{1 + m\mathbf{S}^{(t+1)}_{\text{BF}}} \in [0,1], \tag{S12}$$

where $m > 0$ is a positive free parameter. The parameter $m$ controls the sharpness of the transition.

With this modulated adaptation rate, the change in a participant's belief is given by an update of the Dirichlet parameters $\alpha^{(t+1)}_{\tilde{s},\tilde{a}}(\tilde{s}')$ for all $(\tilde{s}, \tilde{s}', \tilde{a}) \in \mathcal{S} \times \mathcal{S} \times \mathcal{A}$ [14]

$$\alpha^{(t+1)}_{\tilde{s},\tilde{a}}(\tilde{s}') = \begin{cases} (1-\gamma_{t+1})\alpha^{(t)}_{\tilde{s},\tilde{a}}(\tilde{s}') + \gamma_{t+1}\alpha^{(1)}(\tilde{s}') + \delta(s', \tilde{s}') & \text{if} \quad \tilde{s} = s, \tilde{a} = a \\ \alpha^{(t)}_{\tilde{s},\tilde{a}}(\tilde{s}') & \text{otherwise} \end{cases}, \tag{S13}$$

where $\gamma_{t+1} = \gamma(\mathbf{S}^{(t+1)}_{\text{BF}}, m)$. The update rule becomes the same as the one in Eq. 6 of the main text if we replace $\alpha^{(t)}_{\tilde{s},\tilde{a}}(\tilde{s}')$ by $\tilde{C}^{(t)}_{\tilde{s},\tilde{a}\to\tilde{s}'} + \epsilon$. The update rule expresses the new belief as a mix between two possibilities, represented by the current parameters $\alpha^{(t)}_{\tilde{s},\tilde{a}}(\tilde{s}')$ and the prior $\alpha^{(1)}(\tilde{s}')$, weighted with $1 - \gamma_{t+1}$ and $\gamma_{t+1}$, respectively. In the case of a large surprise, the value of $\gamma_{t+1}$ is close to one, and as a result, the current parameters are forgotten. The update makes a step based on the currently observed transition, expressed by the Kronecker-$\delta$ in the first line. The parameters of transitions from the pairs of the states and actions different form the current one (i.e., $s$ and $a$) are not changed (second line). The update rule of Eq. S13 is called Variational Surprise Minimizing Learning (VarSMiLe) rule in [14].

**Formalizing model-based $Q$-values.** The world-model of the participants is summarized by their beliefs $q^{(t)}(\theta)$ about the transition matrix of the environment. The belief is used for evaluation of two sets of $Q$-values [21], one for novelty $N$ and the other one for the external reward $R$.

Novelty $N^{(t)}(s)$ of state $s$ at time $t$ (cf. Eq. S4) is useful to guide behavior during exploration. Analogous to the common framework in reinforcement learning [21] where information of a reward at state $s'$ is propagated by the Bellman equation to states $s \neq s'$, we use a Bellman equation to propagate the novelty of state $s'$ to other states $s \neq s'$ by using the model of the world. More specifically, for the model-based branch, we assign to each state-action pair a novelty-based value

$Q^{(t)}_{\text{MB,N}}(s,a)$ which is an estimation of the accumulated future discounted novelty that can be gained by taking action $a$ in state $s$. The Bellman equation is

$$Q^{(t)}_{\text{MB,N}}(s,a) = \sum_{s' \in \mathcal{S}} \hat{\theta}^{(t)}_{s,a}(s') \Big( N^{(t)}(s') + \lambda_N \max_{a' \in \mathcal{A}} Q^{(t)}_{\text{MB,N}}(s',a') \Big), \tag{S14}$$

where $\hat{\theta}^{(t)}_{s,a}(s')$ are the estimated transition probabilities, and $\lambda_N \in [0,1]$ is a discount factor for novelty. The Bellman equation assigns a value to the action $a$ in state $s$ as long as a novel state is likely to be reached within the next few steps - even if the immediately neighboring states are not novel. The discount rate $\lambda_N$ controls the time horizon of 'future novelty'. For $\lambda_N \to 0$, only the novelty of the immediately following state matters; for $\lambda_N \to 1$, the time horizon becomes infinitely long.

Rewards $R(s)$ of states $s \in \mathcal{S}$ guide behavior during exploitation. In the theory of reinforcement learning, reward information is summarized in values $Q^{(t)}_{\text{MB,R}}(s,a)$ that are estimations of the accumulated future discounted reward that can be collected when starting at state $s$ with action $a$. The $Q$-values are given by the Bellman equation

$$Q^{(t)}_{\text{MB,R}}(s,a) = \sum_{s' \in \mathcal{S}} \hat{\theta}^{(t)}_{s,a}(s') \Big( R(s') + \lambda_R \max_{a' \in \mathcal{A}} Q^{(t)}_{\text{MB,R}}(s',a') \Big), \tag{S15}$$

where $\lambda_R \in [0,1]$ is the discount factor for reward, which is not necessarily equal to the discount factor for novelty $\lambda_N$. Note that in our environment $R(s) = 0$ at all states except at the goal. Since the scale of the reward is arbitrary, we set $R(s_{Goal}) = 1$. As a result, the reward function is $R(s) = \delta(s, s_{Goal})$.

The total model-based Q-value is a linear combination of the $Q$-values for novelty $Q^{(t)}_{\text{MB,N}}(s,a)$ and reward $Q^{(t)}_{\text{MB,N}}(s,a)$,

$$Q^{(t)}_{\text{MB}}(s,a) = Q^{(t)}_{\text{MB,R}}(s,a) + \beta_N Q^{(t)}_{\text{MB,N}}(s,a), \tag{S16}$$

where $\beta_N \geq 0$ is a free parameter controlling the trade-off between exploitation and exploration, i.e., between reward-seeking and novelty-seeking behavior.

In our model, $\beta_N$ depends on whether participants are in the exploration phase or the exploitation phase. This dependency is simplified as follows: Since novelty is the main drive in the 1st episode of the 1st block, we keep $\beta_N$ fixed at a value $\beta_{N1}$ throughout this episode. However, at the end of the 1st episode of the 1st block, once participants have found the goal and do not need further exploration, we set $\beta_N = 0$ and keep it at zero for all remaining episodes of the 1st block.

Surprise increases rapidly at the first mismatch that participants face in the 1st episode of the 2nd block, when they encounter an unexpected transition. We hypothesize that the huge surprise signal triggers renewed exploration and we therefore set $\beta_N = \beta_{N2}$ for the 1st episode of the 2nd block. With the same arguments as for the 1st block, we set $\beta_N$ to zero for the remaining episodes of the 2nd block. $\beta_{N1}$ and $\beta_{N2}$ are free parameters of the model.

Note that, for model comparison, we use the same assumptions for all other alternative algorithms that either seek novelty or uncertainty - see Section 2.

**Updating model-based Q-value.** Since solving the non-linear equations S14 and S15 for computing two separate sets of model-based $Q$-values (i.e., $Q^{(t)}_{\text{MB,N}}(s,a)$ and $Q^{(t)}_{\text{MB,R}}(s,a)$ for all

$(\tilde{s}, \tilde{a}) \in \mathcal{S} \times \mathcal{A})$ is computationally costly, we use a variant (Algorithm 4) of Prioritized Sweeping [21–23].

The idea of the algorithm, for example for updating $Q^{(t)}_{\text{MB,R}}(s, a)$, is to define a set of $|\mathcal{S}|$ mirror variables $U^{(t)}_{\text{R}}(s)$, and rewrite Eq. S15 as

$$\begin{aligned} Q^{(t)}_{\text{MB,R}}(s, a) &= \sum_{s' \in \mathcal{S}} \hat{\theta}^{(t)}_{s,a}(s')\Big(R(s') + \lambda_R U^{(t)}_{\text{R}}(s')\Big) \\ U^{(t)}_{\text{R}}(s') &= \max_{a' \in \mathcal{A}} Q^{(t)}_{\text{MB,R}}(s', a'). \end{aligned} \tag{S17}$$

At the transition from time step $t-1$ to time step $t$ several iterations take place. The algorithm first puts $U^{(t)}_{\text{R}}(s) = U^{(t-1)}_{\text{R}}(s)$ and updates $Q^{(t)}_{\text{MB,R}}(s, a)$ for all $s, a$ with the current values of $U^{(t)}_{\text{R}}(s)$ using the 1st equation. The size of the update step for the value of a state $s'$ is measured as $\Delta V(s') = |U^{(t)}_{\text{R}}(s') - \max_{a' \in \mathcal{A}} Q^{(t)}_{\text{MB,R}}(s', a')|$. The states $s'$ are then ordered in a priority queue with the state of biggest update step at the top. The algorithm updates the values of $U^{(t)}_{\text{R}}(s')$ of the top priority state using the 2nd equation. This results in further updates $\Delta Q^{(t)}_{\text{MB,R}}(s, a) = \hat{\theta}^{(t)}_{s,a}(s')\lambda_R \Delta V(s')$ for all $s, a$ induced by the first equation. After these updates the priority list is resorted. Updating ends after $T_{\text{PS}}$ iterations where $T_{\text{PS}} \in \mathbb{N}$ is a free parameter of the algorithm.

The values of $Q^{(1)}_{\text{MB,R}}(s, a)$, $U^{(1)}_{\text{R}}(s)$, $Q^{(1)}_{\text{MB,N}}(s, a)$, and $U^{(1)}_{\text{N}}(s)$ are initialized consistent with the Bellman equations under the prior world-model (uniform distribution for all transitions) and the prior reward (zero) and novelty values ($\log |\mathcal{S}|$). For details, see Algorithms 1 and 4.

#### 1.2 SurNoR model-free branch

The pseudocode for the model-free branch is shown in Alg. 3. In this subsection, the details are explained.

**Formalizing model-free $Q$-values.** Analogous to the model-based branch, we define $Q^{(t)}_{\text{MF,R}}(s, a)$ and $Q^{(t)}_{\text{MF,N}}(s, a)$ as values of the state-action pairs corresponding to the external reward $R$ and novelty $N$, respectively. In contrast to the model-based branch, the model-free $Q$-values are updated using TD-learning [21, 24], for which the model of the world is not directly used - see the paragraph 'Updating model-free $Q$-values'.

Analogous to the total model-based $Q$-values, we define the total model-free $Q$-values as

$$Q^{(t)}_{\text{MF}}(s, a) = Q^{(t)}_{\text{MF,R}}(s, a) + \beta_N Q^{(t)}_{\text{MF,N}}(s, a), \tag{S18}$$

where $\beta_N \geq 0$ has the same value as the one used in Eq. S16.

**Reward and novelty prediction errors.** A crucial signal in model-free reinforcement learning is the reward prediction error (RPE), defined as the difference between the expected 'reward' of a state-action pair and its real 'reward' [21]. Since we defined two separate sets of $Q$-values, one for the external reward and one for novelty (which plays the role of an 'internal reward'), we also define two separate corresponding prediction errors.

Consider the transition $(S_t = s, A_t = a) \to (S_{t+1} = s')$. The RPE at time $t+1$ is defined as

$$RPE_{t+1} = R(s') + \lambda_R \max_{a' \in \mathcal{A}} Q^{(t)}_{\text{MF,R}}(s', a') - Q^{(t)}_{\text{MF,R}}(s, a), \tag{S19}$$

and similarly, the novelty prediction error (NPE) at time $t+1$ is defined as

$$NPE_{t+1} = N^{(t)}(s') + \lambda_N \max_{a' \in \mathcal{A}} Q^{(t)}_{\mathrm{MF,N}}(s', a') - Q^{(t)}_{\mathrm{MF,N}}(s, a), \tag{S20}$$

where $\lambda_R$ and $\lambda_N$ are the same discount factors as the ones used in the model-based branch.

**Eligibility trace.** To keep track of the previously chosen state-action pairs, and to include them in the update rule, we use eligibility traces [21, 25, 26]. To have the most general setting, we define two separate eligibility traces, one for the external reward $e^{(t)}_R(s, a)$ and one for novelty (the internal reward) $e^{(t)}_N(s, a)$ for all state-action pairs $(s, a)$. We initialize the eligibility traces at zero and reset their values to zero at the beginning of each episode. After the transition $(S_t = s, A_t = a) \to (S_{t+1} = s')$, the eligibility traces are updated to

$$\begin{aligned} e^{(t+1)}_R(s', a') &= \begin{cases} 1 & \text{if} \quad s' = s, a' = a \\ \lambda_R \mu_R e^{(t)}_R(s', a') & \text{otherwise} \end{cases} \\ e^{(t+1)}_N(s', a') &= \begin{cases} 1 & \text{if} \quad s' = s, a' = a \\ \lambda_N \mu_N e^{(t)}_N(s', a') & \text{otherwise}\,, \end{cases} \end{aligned} \tag{S21}$$

where $\lambda_R$ and $\lambda_N$ are the discount factors defined above, and $\mu_N \in [0, 1]$ and $\mu_R \in [0, 1]$ are free parameters expressing how fast eligibility traces decay in time.

**Surprise modulation of model-free learning rate.** Usual TD learning algorithms use a constant (or decreasing in time) learning rate for updating $Q$-values [21]. However, the model-free branch of SurNoR has a learning rate modulated by the model-based branch. This novel interaction between model-based and model-free modules has not been explored by previous hybrid models in neuroscience, e.g., [1, 2].

We define the surprise modulated model-free learning rate $\rho_t$ as

$$\rho_t = \rho_b + \gamma(\mathbf{S}^{(t)}_{\mathrm{BF}}, m)\delta\rho, \tag{S22}$$

where $\gamma(\mathbf{S}^{(t)}_{\mathrm{BF}}, m)$ is the surprise modulated adaptation rate of the model-based branch defined in Eq. S12, $\rho_b \in [0, 1]$ is the baseline learning rate (when there is no surprise, i.e., if $\mathbf{S}^{(t)}_{\mathrm{BF}} = 0$), and $\delta\rho \in [0, 1 - \rho_b]$ is the maximum possible variation of the learning rate due to the surprise modulation. As a result, the learning rate value $\rho_t$ ranges between $\rho_b$ (when $\mathbf{S}^{(t)}_{\mathrm{BF}} = 0$) and $\rho_b + \delta\rho$ (when $\mathbf{S}^{(t)}_{\mathrm{BF}} \to \infty$).

**Updating model-free Q-value.** The model-free $Q$-values for external reward are initialized to zero, $Q^{(1)}_{\mathrm{MF,R}}(s, a) = 0$ for all $s, a$. This initialization avoids any potential bias towards optimistic initialization (OI). The reason for this choice is to have novelty as the only exploration drive during the 1st episode of the 1st block. We separately test the effect of the initialization of reward-based $Q$-values in three alternative algorithm which use OI of $Q^{(1)}_{\mathrm{MF,R}}(s, a)$ as a drive for exploration [21] - see Section 2. However, to consider the most general case, we initialize the model-free $Q$-values for novelty at $Q^{(1)}_{\mathrm{MF,N}}(s, a) = Q_{N0}$ with a free parameter $Q_{N0} \geq 0$.

At each time step $t+1$, the model-free $Q$-values are updated with a TD-learning algorithm

$$\begin{aligned} Q_{\mathrm{MF,R}}^{(t+1)}(s,a) &= Q_{\mathrm{MF,R}}^{(t)}(s,a) + \rho_{t+1} e_R^{(t+1)}(s,a) RPE_{t+1} \\ Q_{\mathrm{MF,N}}^{(t+1)}(s,a) &= Q_{\mathrm{MF,N}}^{(t)}(s,a) + \rho_{t+1} e_N^{(t+1)}(s,a) NPE_{t+1}. \end{aligned} \tag{S23}$$

for all $(s,a) \in \mathcal{S} \times \mathcal{A}$.

#### 1.3 Hybrid policy

The policy for action selection is based on a linear combination of $Q$-values, similar to [1, 2]. We use a softmax policy [21] and consider the probability of choosing action $a$ in state $s$ as

$$\pi(A_t = a | S_t = s) = \frac{1}{Z(s)} \exp\left\{ \beta \left[ \omega \left( \omega_{\mathrm{scale}} Q_{\mathrm{MF}}^{(t)}(s,a) \right) + (1-\omega)\, Q_{\mathrm{MB}}^{(t)}(s,a) \right] \right\}, \tag{S24}$$

where $Z(s)$ is the normalization constant (that ensures that $\sum_a \pi(A_t = a|S_t = s) = 1$), $\omega_{\mathrm{scale}} \geq 0$ is a free parameter to correct the potentially different scaling of the model-based and model-free values, and $\omega \in [0,1]$ is a free parameter to balance the relative contribution of the model-based and model-free branches on the policy. When $\omega = 1$, the policy is purely model-free (but includes the effect of surprise modulation on the TD-learning learning rate), and when $\omega = 0$, the policy is purely model-based. Note that $\omega_{MF}$ and $\omega_{MB}$ mentioned in the main texts are equal to $\omega \times \omega_{\mathrm{scale}}$ and $1-\omega$, respectively. The reverse temperature $\beta \geq 0$ controls the sharpness of policy (the larger $\beta$ the more deterministic is the policy).

As it was shown by [1], $\omega$ can vary in time. Specific to our experiment, we consider $\omega$ to be piece-wise constant in time: 1. $\omega = \omega_{11}$ for the 1st episode of the 1st block, when participants are in the pure exploration phase, 2. $\omega = \omega_{12}$ for the 1st episode of the 2nd block, when the goal is lost, and 3. $\omega = \omega_0$ for the rest of the experiments (i.e., episodes 2 to 5 for both blocks), when participants are in the exploitation phase. Moreover, we allow the value of $\beta$ to be different for the 1st and the 2nd block, $\beta_1$ and $\beta_2$ respectively. By doing so, we allow the model to change its confidence in action selection after observing the sudden change in the environment.

Note that, for model comparison, we use the same assumptions for all other alternative algorithms that use hybrid policy - see Section 2.

#### 1.4 Summary of free parameters

SurNoR has 18 free parameters, summarized as

$$\{\epsilon, m, \lambda_R, \lambda_N, \beta_1, \beta_2, \beta_{N1}, \beta_{N2}, T_{\mathrm{PS}}, \mu_R, \mu_N, Q_{N0}, \rho_b, \delta\rho, \omega_{\mathrm{scale}}, \omega_0, \omega_{11}, \omega_{12}\}. \tag{S25}$$

$\epsilon$ is used for initialization of the belief in Eq. S9. $m$ is used for modulation of the adaptation rate in Eq. S12. $\lambda_R$ and $\lambda_N$ are discount factors used in the definitions and the updates of $Q$-values. $\beta_1$ and $\beta_2$ are the inverse temperatures controlling the sharpness of the hybrid policy in Eq. S24. $\beta_{N1}$ and $\beta_{N2}$ are used for balancing novelty against external reward in equations S16 and S18. $T_{\mathrm{PS}}$ is used for Prioritized Sweeping in Algorithm 4. $\mu_R$ and $\mu_N$ are used for controlling the decay of eligibility traces in Eq. S21. $Q_{N0}$ is used for initialization of $Q_{\mathrm{MF,N}}$. $\rho_b$ and $\delta\rho$ are used for the baseline learning rate of the model-free branch and its surprise modulation in Eq. S22. $\omega_{\mathrm{scale}}$ is used for correcting the potential different scaling of the model-based and model-free values in Eq. S24, and $\omega_0$, $\omega_{11}$, and $\omega_{12}$ are used for balancing model-free against model-based in the hybrid policy of Eq. S24.

**Algorithm 1** Pseudocode for SurNoR

Specify $\mathcal{S}$ and $\mathcal{A}$<br>
Specify Episode (Epi) and Block<br>
### Parameter specification<br>
Specify $\{\epsilon, m, \lambda_R, \lambda_N, \beta_1, \beta_2, \beta_{N1}, \beta_{N2}, T_{\text{PS}}, \mu_R, \mu_N, Q_{N0}, \rho_b, \delta\rho, \omega_{\text{scale}}, \omega_0, \omega_{11}, \omega_{12}\}$.<br>
**if** Block $= 1$ and Epi $= 1$, **then** Put $\beta = \beta_1$, $\omega = \omega_{11}$ and $\beta_N = \beta_{N1}$.<br>
**if** Block $= 1$ and Epi $\neq 1$, **then** Put $\beta = \beta_1$, $\omega = \omega_0$ and $\beta_N = 0$.<br>
**if** Block $= 2$ and Epi $= 1$, **then** Put $\beta = \beta_2$, $\omega = \omega_{12}$ and $\beta_N = \beta_{N2}$.<br>
**if** Block $= 2$ and Epi $\neq 1$, **then** Put $\beta = \beta_2$, $\omega = \omega_0$ and $\beta_N = 0$.<br>
### Initialization<br>
Put $e_R^{(1)}(s,a) = e_N^{(1)}(s,a) = 0$, $\forall (s,a) \in \mathcal{S} \times \mathcal{A}$.<br>
**if** Epi $= 1$ and Block $= 1$ **then**<br>
    Put $C_s^{(1)} = 0$, $U_R^{(1)}(s) = 0$, $U_N^{(1)}(s) = \frac{\log(|\mathcal{A}|)}{1-\lambda}$, $\forall s \in \mathcal{S}$.<br>
    Put $Q_{\text{MB,R}}^{(1)}(s,a) = 0$, $Q_{\text{MB,N}}^{(1)}(s,a) = U_N^{(1)}(s)$, $\forall (s,a) \in \mathcal{S} \times \mathcal{A}$.<br>
    Put $Q_{\text{MF,R}}^{(1)}(s,a) = 0$, $Q_{\text{MF,N}}^{(1)}(s,a) = Q_{N0}$, $\forall (s,a) \in \mathcal{S} \times \mathcal{A}$.<br>
    Put $\alpha_{s,a}^{(1)}(s') = \epsilon$, $\forall (s,s',a) \in \mathcal{S} \times \mathcal{S} \times \mathcal{A}$.<br>
**else**<br>
    Initialize $C_s^{(1)}$, $U_R^{(1)}(s)$, $U_N^{(1)}(s)$, $Q_{\text{MB,R}}^{(1)}(s,a)$, $Q_{\text{MB,N}}^{(1)}(s,a)$, $Q_{\text{MF,R}}^{(1)}(s,a)$, $Q_{\text{MF,N}}^{(1)}(s,a)$ and $\alpha_{s,a}^{(1)}(s')$ with their latest values in the previous Episode.<br>
Initialize state $S_1 = s_1$ and update counts $C_s^{(1)} \leftarrow C_s^{(1)} + \delta(s, s_1)$.<br>
$t \leftarrow 1$.<br>
### Going through the task<br>
**while** $s_t \neq s_{Goal}$ **do**<br>
### Making action<br>
    Compute $Q_{\text{MF}}^{(t)}(s,a) = Q_{\text{MF,R}}^{(t)}(s,a) + \beta_N Q_{\text{MF,N}}^{(t)}(s,a)$.<br>
    Compute $Q_{\text{MB}}^{(t)}(s,a) = Q_{\text{MB,R}}^{(t)}(s,a) + \beta_N Q_{\text{MB,N}}^{(t)}(s,a)$.<br>
    Sample $a_t$ from $\pi(A_t = a | S_t = s) \propto \exp\Big\{\beta\Big[\omega\left(\omega_{\text{scale}} Q_{\text{MF}}^{(t)}(s,a)\right) + (1-\omega)\, Q_{\text{MB}}^{(t)}(s,a)\Big]\Big\}$.<br>
    Observe $S_{t+1} = s_{t+1}$.<br>
### Updating internal variables<br>
    Update counts $C_s^{(t+1)} = C_s^{(t)} + \delta(s, s_{t+1})$ and novelty $N^{(t+1)}(s) = \log \frac{t + |\mathcal{S}|}{C_s^{(t+1)} + 1}$.<br>
    Update $\alpha^{(t+1)}$, $U_R^{(t+1)}$, $U_N^{(t+1)}$, $Q_{\text{MB,R}}^{(t+1)}$, and $Q_{\text{MB,N}}^{(t+1)}$ using the model-based branch in Alg. 2.<br>
    Update $e_N^{(t+1)}$, $e_R^{(t+1)}$, $Q_{\text{MF,R}}^{(t+1)}$, and $Q_{\text{MF,N}}^{(t+1)}$ using the model-free branch in Alg. 3.<br>
### Going to the next step<br>
    $t \leftarrow t + 1$.

**Algorithm 2** Pseudocode for the model-based branch of SurNoR

```
# Surprise and adaptation rate
```
Compute $\mathbf{S}^{(t+1)} = \hat{\theta}^{(1)}_{s_t,a_t}(s_{t+1})/\hat{\theta}^{(t)}_{s_t,a_t}(s_{t+1})$.

Compute $\gamma_{t+1} = m\mathbf{S}^{(t+1)}/\big(1 + m\mathbf{S}^{(t+1)}\big)$.

```
# Updating the belief
```
Update $\alpha^{(t+1)}_{s_t,a_t}(s) = (1-\gamma_{t+1})\alpha^{(t)}_{s_t,a_t}(s) + \gamma_{t+1}\epsilon + \delta(s_{t+1}, s)$, $\forall s \in \mathcal{S}$.

Update $\alpha^{(t+1)}_{s,a}(s') = \alpha^{(t)}_{s,a}(s')$, $\forall s \neq s_t, a \neq a_t$, and $s' \in \mathcal{S}$.

Update $\hat{\theta}^{(t+1)}$ as $\hat{\theta}^{(t+1)}_{s,a}(s') = \alpha^{(t+1)}_{s,a}(s')/\sum_{\tilde{s}'\in\mathcal{S}} \alpha^{(t+1)}_{s,a}(\tilde{s}')$.

```
# Updating the values
```
Update $Q^{(t+1)}_{\text{MB,N}}(s,a)$ and $U^{(t+1)}_N(s)$ using Alg. 4 and $N^{(t+1)}(s)$ as rewards.

**if** Epi $= 1$ and Block $= 1$ and $s_t \neq s_{Goal}$ **then**

  Update $Q^{(t+1)}_{\text{MB,R}}(s,a) = U^{(t+1)}_R(s) = 0$.

**else**

  Update $Q^{(t+1)}_{\text{MB,R}}(s,a)$ and $U^{(t+1)}_R(s)$ using Alg. 4 and $R(s) = \delta(s, s_{Goal})$ as rewards.

**Algorithm 3** Pseudocode for the model-free branch of SurNoR

```
# Surprise-modulated learning rate
```
Compute $\rho_{t+1} = \rho_b + \gamma_{t+1}\delta\rho$.

```
# Prediction errors
```
Compute $RPE_{t+1} = R(s_{t+1}) + \lambda_R \max_{a'\in\mathcal{A}} Q^{(t)}_{\text{MF,R}}(s_{t+1}, a') - Q^{(t)}_{\text{MF,R}}(s_t, a_t)$.

Compute $NPE_{t+1} = N^{(t)}(s_{t+1}) + \lambda_N \max_{a'\in\mathcal{A}} Q^{(t)}_{\text{MF,N}}(s_{t+1}, a') - Q^{(t)}_{\text{MF,N}}(s_t, a_t)$.

```
# Update of the eligibility traces
```
Update $e^{(t+1)}_N(s_t, a_t) = 1$, and $e^{(t+1)}_N(s,a) = \lambda_N \mu_N e^{(t)}_N(s,a)$, $\forall s \neq s_t, a \neq a_t$.

Update $e^{(t+1)}_R(s_t, a_t) = 1$, and $e^{(t+1)}_R(s,a) = \lambda_R \mu_R e^{(t)}_R(s,a)$, $\forall s \neq s_t, a \neq a_t$.

```
# TD-learners
```
Update $Q^{(t+1)}_{\text{MF,R}}(s,a) = Q^{(t)}_{\text{MF,R}}(s,a) + \rho_{t+1} e^{(t+1)}_R(s,a) RPE_{t+1}$, $\forall s \in \mathcal{S}$ and $a \in \mathcal{A}$.

Update $Q^{(t+1)}_{\text{MF,N}}(s,a) = Q^{(t)}_{\text{MF,N}}(s,a) + \rho_{t+1} e^{(t+1)}_N(s,a) NPE_{t+1}$, $\forall s \in \mathcal{S}$ and $a \in \mathcal{A}$.

**Algorithm 4** Pseudocode for the modified version of Prioritized Sweeping Algorithm for one time-step at time $t+1$

### Specifying whether the update is for the internal or the external reward

Put $\lambda = \lambda_R$ for reward and $\lambda = \lambda_N$ for novelty.

Put $Q^{(t)} = Q^{(t)}_{\text{MB,R}}$, $U^{(t)} = U^{(t)}_R$, and Reward $= R$ for reward, and put $Q^{(t)} = Q^{(t)}_{\text{MB,N}}$, $U^{(t)} = U^{(t)}_N$, and Reward $= N^{(t+1)}$ for novelty.

### Applying the effect of the latest observation on $Q$-values using previous $U$-values

**for** $(s,a) \in \mathcal{S} \times \mathcal{A}$ **do**

  $Q^{(t+1)}(s,a) = \sum_{s' \in \mathcal{S}} \hat{\theta}^{(t+1)}_{s,a}(s') \Big( \text{Reward}(s') + \lambda U^{(t)}(s') \Big)$

### Making the priority queue

**for** $s \in \mathcal{S}$ **do**

  $U^{(t+1)}(s) = U^{(t)}(s)$

  $\text{Prior}(s) = |U^{(t+1)}(s) - \max_{a \in \mathcal{A}} Q^{(t+1)}(s,a)|$

### Updating $U$-values for $T_{\text{PS}}$ steps

**for** $T_{\text{PS}}$ iterations **do**

  $s' = \arg\max_{s \in \mathcal{S}} \text{Prior}(s)$

  $\Delta V = \max_{a \in \mathcal{A}} Q^{(t+1)}(s',a) - U^{(t+1)}(s')$

  $U^{(t+1)}(s') = \max_{a \in \mathcal{A}} Q^{(t+1)}(s',a)$

  # Applying the effect of the update of $U$-values on $Q$-values

  **for** $(s,a) \in \mathcal{S} \times \mathcal{A}$ **do**

    $Q^{(t+1)}(s,a) \leftarrow Q^{(t+1)}(s,a) + \lambda \hat{\theta}^{(t+1)}_{s,a}(s') \Delta V$

  # Updating the priority queue

  **for** $s \in \mathcal{S}$ **do**

    $\text{Prior}(s) = |U^{(t+1)}(s) - \max_{a \in \mathcal{A}} Q^{(t+1)}(s,a)|$

### 2 Alternative algorithms

To statistically test the effect of surprise and novelty, we implemented 12 alternative algorithms explained in this section. Their key features are summarized in Table 1. The modified versions of SurNoR which do not seek novelty but assign negative reward to the most frequent states are explained at the end.

**Model-based alternatives.** Four out of 12 algorithms are purely model-based. They all use world-model and prioritized sweeping to calculate model-based Q-values. However, they have different approaches for learning the world-model and different strategies for exploration.

(i) MB+S+N: This algorithm has both features of SurNoR in using surprise modulation for model-building and novelty-seeking for exploration, but it does not use a parallel TD-learner. MB+S+N is a reduced version of SurNoR with $\mu_R = \mu_N = Q_{N0} = \rho_b = \delta\rho = \omega_{\text{scale}} = \omega_0 = \omega_{11} = \omega_{12} = 0$, which is equivalent to the model-based branch of SurNoR. MB+S+N has 9 free parameters $\{\epsilon, m, \lambda_R, \lambda_N, T_{PS}, \beta_1, \beta_2, \beta_{N1}, \beta_{N2}\}$.

(ii) MB+N: This algorithm is a modified version of MB+S+N; it uses novelty-seeking for exploration, but it does not have surprise modulation for learning the world-model. MB+N uses leaky integration to update the belief parameters (analogous to Eq. S13),

$$\alpha^{(t+1)}_{\tilde{s},\tilde{a}}(\tilde{s}') = \begin{cases} \kappa_{\text{Leak}}\alpha^{(t)}_{\tilde{s},\tilde{a}}(\tilde{s}') + \delta(s', \tilde{s}') & \text{if} \quad \tilde{s} = s, \tilde{a} = a \\ \alpha^{(t)}_{\tilde{s},\tilde{a}}(\tilde{s}') & \text{otherwise} \end{cases}, \tag{S26}$$

where $\kappa_{\text{Leak}} \in [0, 1]$ is a constant free parameter. Such a learning rule has been used previously to model human behavior [27–30]. Overall, MB+N has 9 free parameters $\{\epsilon, \kappa_{\text{Leak}}, \lambda_R, \lambda_N, T_{PS}, \beta_1, \beta_2, \beta_{N1}, \beta_{N2}\}$. It cannot be considered as a special case of SurNoR, but it can be implemented in the framework of the SurNoR algorithm by using Eq. S26 instead of Eq. S13 for updating the belief and by putting $m = \mu_R = \mu_N = Q_{N0} = \rho_b = \delta\rho = \omega_{\text{scale}} = \omega_{11} = \omega_{12} = \omega_0 = 0$.

(iii) MB+S+U: This algorithm is similar to MB+S+N; it uses surprise modulation for learning the world-model, but it seeks uncertainty instead of novelty for exploration. Following the ideas from [8, 9], we define a set of uncertainty-based Q-values, analogous to the SurNoR's novelty-based Q-values (Eq. S14), as

$$Q^{(t)}_{\text{MB,U}}(s, a) = \sum_{s' \in \mathcal{S}} \hat{\theta}^{(t)}_{s,a}(s')\Big( -\log \hat{\theta}^{(t)}_{s,a}(s') + \lambda_U \max_{a' \in \mathcal{A}} Q^{(t)}_{\text{MB,U}}(s', a')\Big), \tag{S27}$$

where $-\log \hat{\theta}^{(t)}_{s,a}(s')$, sometimes called surprisal (equal to Shannon surprise) is considered as the intrinsic reward of the transition $(s, a) \to s'$. The model MB+S+U is implemented by modifying SurNoR in 3 steps: 1. Replacing Eq. S14 by Eq. S27 and using $Q^{(t)}_{\text{MB,U}}(s, a)$ instead of $Q^{(t)}_{\text{MB,N}}(s, a)$ in all equations. 2. Replacing $\lambda_N$ by $\lambda_U$, $\beta_{N1}$ by $\beta_{U1}$, and $\beta_{N2}$ by $\beta_{U2}$. 3. Putting $\mu_R = \mu_N = Q_{N0} = \rho_b = \delta\rho = \omega_{\text{scale}} = \omega_0 = \omega_{11} = \omega_{12} = 0$. MB+S+U has 9 free parameters $\{\epsilon, m, \lambda_R, \lambda_U, T_{PS}, \beta_1, \beta_2, \beta_{U1}, \beta_{U2}\}$.

(iv) MB+S+OI: This algorithm removes the novelty-seeking block of MB+S+N and uses optimistic initialization for exploration, i.e., it updates reward-based Q-values also in the 1st episode of block 1 even before observing the goal states. MB+S+OI can be implemented by modifying SurNoR in 2 steps: 1. Removing the 'if' condition in the lines 7-10 of Alg. 2 and keeping only line 10. 2.

Putting $\lambda_N = \beta_{N1} = \beta_{N2} = \mu_R = \mu_N = Q_{N0} = \rho_b = \delta\rho = \omega_{\text{scale}} = \omega_0 = \omega_{11} = \omega_{12} = 0$. This algorithm has 6 free parameters $\{\epsilon, m, \lambda_R, T_{PS}, \beta_1, \beta_2\}$.

**Model-free alternatives.** Four out of 12 algorithms are model-free. All of them use a TD-learner for learning model-free Q-values. However, the ones with surprise-modulation are also equipped with a world-model, but the world model is not used for computing a set of model-based Q-values, and the policy is not hybrid.

(v) MF+S+N: This algorithm is equivalent to the model-free branch of SurNoR, but it also uses the world-model in the model-based branch for surprise-computation. MF+S+N can be seen as a reduced version of SurNoR by putting $T_{PS} = 0$ and $\omega_{\text{scale}} = \omega_{11} = \omega_{12} = \omega_0 = 1$. It has 13 free parameters $\{\epsilon, m, \lambda_R, \lambda_N, \beta_1, \beta_2, \beta_{N1}, \beta_{N2}, \mu_R, \mu_N, Q_{N0}, \rho_b, \delta\rho\}$.

(vi) MF+N: This algorithm is a reduced version of SurNoR by putting $m = T_{PS} = \delta\rho = 0$ and $\omega_{\text{scale}} = \omega_{11} = \omega_{12} = \omega_0 = \epsilon = 1$, which is equivalent to the model-free branch of SurNoR without any surprise modulation. It can also be seen as a modified version of the famous $Q(\lambda)$ algorithm [21] (with $\lambda = \mu_R$ in our notation) but with novelty as an exploration bonus (instead of optimistic initialization). The model MF+N has overall 10 free parameters $\{\lambda_R, \lambda_N, \beta_1, \beta_2, \beta_{N1}, \beta_{N2}, \mu_R, \mu_N, Q_{N0}, \rho_b\}$.

(vii) MF+S+U: The relation between MF+S+U and MB+S+U is the same as the relation between MF+S+N and MB+S+N. All the features of MF+S+U (including the surprise modulation of the learning rate of the model-free) except for its exploration strategy are the same as the ones of MF+S+N. For exploration, MF+S+U seeks uncertainty instead of novelty. Similar to what we did for MB+S+U, we followed the ideas from [8, 9] and defined the uncertainty-based Q-values as in Eq. S27. Then, we define the Uncertainty Prediction Error (UPE), analogous to the SurNoR's NPE (Eq. S20), as

$$UPE_{t+1} = -\log \hat{\theta}^{(t)}_{s,a}(s') + \lambda_U \max_{a' \in \mathcal{A}} Q^{(t)}_{\text{MF,U}}(s', a') - Q^{(t)}_{\text{MF,U}}(s, a), \tag{S28}$$

and then we update the uncertainty-based model-free Q-values as

$$Q^{(t+1)}_{\text{MF,U}}(s, a) = Q^{(t)}_{\text{MF,U}}(s, a) + \rho_{t+1} e^{(t+1)}_U(s, a) UPE_{t+1}, \tag{S29}$$

where $e^{(t+1)}_U(s, a)$ is the uncertainty eligibility trace with a decay factor $\mu_U$. Then MF+S+U can be implemented by modifying SurNoR in three steps: 1. Replacing $Q^{(t)}_{\text{MF,N}}(s, a)$ by $Q^{(t)}_{\text{MF,U}}(s, a)$ in all equations. 2. Replacing $\lambda_N$ by $\lambda_U$, $\beta_{N1}$ by $\beta_{U1}$, $\beta_{N2}$ by $\beta_{U2}$, and $\mu_N$ by $\mu_U$. 3. Putting $T_{PS} = 0$ and $\omega_{\text{scale}} = \omega_{11} = \omega_{12} = \omega_0 = 1$. The model MF+S+U has 13 free parameters $\{\epsilon, m, \lambda_R, \lambda_U, \beta_1, \beta_2, \beta_{U1}, \beta_{U2}, \mu_R, \mu_U, Q_{U0}, \rho_b, \delta\rho\}$.

(iix) MF+OI: This algorithm is our simplest algorithm, and neither surprise nor novelty is used in it. MF+OI is equivalent to $Q(\lambda)$[21], with $\lambda = \mu_R$ in our notation. It uses optimistic initialization for exploration by putting $Q^{(0)}_{MF,R} = Q_{R0}$, where $Q_{R0}$ is a free parameter. It can be seen as a modified version of SurNoR by initializing $Q^{(0)}_{MF,R} = Q_{R0}$ and putting $m = \lambda_N = \beta_{N1} = \beta_{N2} = T_{PS} = \mu_N = Q_{N0} = \delta\rho = 0$ and $\omega_{\text{scale}} = \omega_{11} = \omega_{12} = \omega_0 = \epsilon = 1$. The model MF+OI has overall 6 free parameters $\{\lambda_R, Q_{R0}, \rho_b, \beta_1, \beta_2, \mu_R\}$.

**Hybrid alternatives.** Three out of 12 algorithms are hybrid, meaning they use both model-free and model-based Q-values for decision-making.

(ix) Hyb+N: This algorithm uses MB+N and MF+N in parallel and combines their Q-values (in the same fashion as in SurNoR) in a hybrid policy. It has overall 17 free parameters $\{\epsilon, \kappa_{\text{Leak}}, \lambda_R, \lambda_N, \beta_1, \beta_2, \beta_{N1}, \beta_{N2}, T_{PS}, \mu_R, \mu_N, Q_{N0}, \rho_b, \omega_{\text{scale}}, \omega_{11}, \omega_{12}, \omega_0\}$.

(x) Hyb+S+U: This algorithm uses MB+S+U and MF+S+U in parallel and combines their Q-values (in the same fashion as in SurNoR) in a hybrid policy. Hyb+S+U is as complex as SurNoR and has overall 18 free parameters $\{\epsilon, m, \lambda_R, \lambda_U, \beta_1, \beta_2, \beta_{U1}, \beta_{U2}, T_{PS}, \mu_R, \mu_U, Q_{U0}, \rho_b, \delta\rho, \omega_{\text{scale}}, \omega_{11}, \omega_{12}, \omega_0\}$.

(xi) Hyb+S+OI: This algorithm uses MF+OI (but with surprise modulation of the learning rate of the model-free branch) and MB+S+OI in parallel and combines their Q-values (in the same fashion as in SurNoR) in a hybrid policy. Hyb+S+OI has overall 14 free parameters $\{\epsilon, m, \lambda_R, \beta_1, \beta_2, T_{PS}, \mu_R, Q_{R0}, \rho_b, \delta\rho, \omega_{\text{scale}}, \omega_{11}, \omega_{12}, \omega_0\}$.

**Null model.** (xii) RC (Random Choice): According to this algorithm, participants choose actions with uniform distribution, i.e., each action is selected with a probability equal to $\frac{1}{|\mathcal{A}|} = 0.25$. We used this model as a reference to quantify the effect of our novelty-seeking exploration in the 1st episode of the 1st block. This algorithm does not have any free parameter.

**Control modifications of SurNoR.** (xiii) Binary Novelty: The two control algorithms mentioned in the main text are exactly the same as SurNoR except for a change in the intrinsic motivation signal that drives exploration. While in the SurNoR algorithm the continuous-valued novelty signal defined in Eq. S3 and Eq. S4 serves as the intrinsic reward, in the control algorithms the intrinsic reward of state $s$ at time $t$ is binary: in the first control algorithm it is considered to be $-1$ if the count $C_s^{(t)} \geq C_{\text{thr}}$ and 0 otherwise, where $C_{\text{thr}}$ is a new free parameter, i.e., the algorithm considers the states that are encountered more than $C_{\text{thr}}$ times as bad states and assigns a constant negative reward to them. Similarly, in the 2nd control algorithm, the intrinsic reward of state $s$ at time $t$ is considered to be $-1$ if state $s$ is among the $n$ most frequently encountered states and 0 otherwise, where $n$ is a new free parameter, i.e., the algorithm considers the $n$ most frequently encountered states as bad states. Therefore, the pseudo-code of the control algorithms is the same as the pseudo-code of SurNoR in Alg. 1 but with 2 modifications: (i) $U_N^{(1)}(s)$ is initialized at a value 0. (ii) The definition of novelty is changed to

$$N^{(t)}(s) = \begin{cases} -1 & \text{if} \quad C_s^{(t)} \geq C_{\text{thr}} \\ 0 & \text{otherwise} \end{cases} \tag{S30}$$

for the 1st algorithm and to

$$N^{(t)}(s) = \begin{cases} -1 & \text{if} \quad C_s^{(t)} \in n \,\text{highest counts} \\ 0 & \text{otherwise} \end{cases} \tag{S31}$$

for the 2nd algorithm. Overall, both algorithm have 19 free parameters, i.e., 18 free parameters of SurNoR plus $C_{\text{thr}}$ for the 1st and $n$ for the 2nd control algorithm.

| | Algorithm | World-model | Hybrid-policy | Novelty | Surprise | Param. |
|---|---|---|---|---|---|---|
| i | MB+S+N | ✓ | ✗ | ✓ | ✓ | 9 |
| ii | MB+N | ✓ | ✗ | ✓ | ✗ | 9 |
| iii | MB+S+U | ✓ | ✗ | ✗ | ✓ | 9 |
| iv | MB+S+OI | ✓ | ✗ | ✗ | ✓ | 6 |
| v | MF+S+N | ✓ | ✗ | ✓ | ✓ | 13 |
| vi | MF+N | ✗ | ✗ | ✓ | ✗ | 10 |
| vii | MF+S+U | ✓ | ✗ | ✗ | ✓ | 13 |
| iix | MF+OI | ✗ | ✗ | ✗ | ✗ | 6 |
| ix | Hyb+N | ✓ | ✓ | ✓ | ✗ | 17 |
| x | Hyb+S+U | ✓ | ✓ | ✗ | ✓ | 18 |
| xi | Hyb+S+OI | ✓ | ✓ | ✗ | ✓ | 14 |
| xii | RC | ✗ | ✗ | ✗ | ✗ | 0 |
| xiii | BinaryNovelty | ✓ | ✓ | (✓) | ✓ | 19 |
| xiv | **SurNoR** | ✓ | ✓ | ✓ | ✓ | 18 |

Table 1: Summary of the key features of all models.

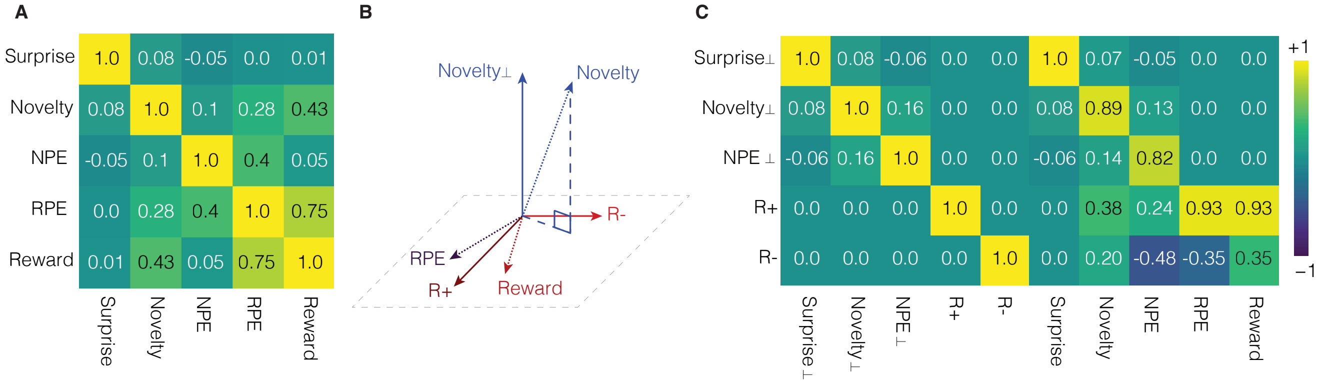


Figure S1: Correlations (averaged over participants) between relevant variables. **A.** The cross-correlations between Surprise, Novelty, NPE, RPE, and Reward during the behavioral task. **B.** Novelty$_\perp$ is the projection of Novelty onto the subspace orthogonal to the plane spanned by Reward and RPE. The variables $R_+$ and $R_-$ are the (normalized) sum and difference of RPE and Reward, respectively. An analogous orthogonalization is applied to Surprise and NPE. **C.** The cross-correlation matrix of the orthogonalized variables and the original ones. Surprise$_\perp$, Novelty$_\perp$, and NPE$_\perp$ are highly correlated with their raw values but have zero correlation with reward and RPE.

### 3 PCA over Reward and RPE

We show that PCA on the two normalized variables Reward and RPE yields $R_+$ and $R_-$.

**Lemma.** For two random variables $X_1$ and $X_2$ with zero mean (i.e., $\mathbb{E}(X_1) = \mathbb{E}(X_2) = 0$), unit variance (i.e., $\mathbb{E}(X_1^2) = \mathbb{E}(X_2^2) = 1$), and correlation $r = \mathbb{E}(X_1 X_2)$, the new variables $X_+ = (X_1 + X_2)/\sqrt{2}$ and $X_- = (X_1 - X_2)/\sqrt{2}$ are the projections on the two normalized principal components of the correlation matrix.

*Proof:* The 2x2 correlation matrix $C$ has diagonal elements $c_{11} = c_{22} = 1$ because of the normalization of each variable to unit variance and off-diagonal elements $c_{21} = c_{12} = r$ according to the assumption and symmetry of correlations. The normalized eigenvectors of the correlation matrix are then $e_+ = (1,1)^T/\sqrt{2}$ and $e_- = (1,-1)^T/\sqrt{2}$ with eigenvalues $\lambda_\pm = 1 \pm r$. ■

Therefore, if Reward and RPE are normalized, then $R_+ = \text{Reward}+\text{RPE}$ and $R_- = \text{Reward}-\text{RPE}$ are their principal components. Furthermore, for positive correlations $r > 0$, the first principal component is $R_+$.

### 4 Correlation and orthogonalization for EEG analysis

Novelty is nearly decorrelated from Surprise and NPE, but Reward and RPE are highly correlated with each other and also correlated with Novelty and NPE (Fig. S1A). PCA on the variables Reward and RPE yields new decorrelated variables $R_+$ and $R_-$ (Section 3). After projecting Surprise, Novelty, and NPE on the space orthogonal to $R_+$ and $R_-$ (Fig. S1B), all five variables are decorrelated (Fig. S1C, left part). Nevertheless, the new variables Surprise$_\perp$, Novelty$_\perp$, and NPE$_\perp$ remain very similar to the original variables Surprise, Novelty, and NPE as indicated by correlations equal to or above 0.89 (Fig. S1C, right part).

### 5 The analysis of random exploration

For any stationary policy (e.g., random choice), the sequence of states $\{S_1, S_2, ...\}$ forms a stationary Markov chain. Let us define the random variable $\mathcal{T}$ as the time of the 1st encounter of the goal state, i.e., $\mathcal{T}$ is the length of the 1st episode. We connect the expected number $\tau_i$ of actions to find the goal starting from state $S_1 = i$ (with $i \in \{1, ..., 10\}$) to the expected number of actions $\tau_j$ in the possible *next* states $S_2 = j$,

$$\tau_i = \mathbb{E}[\mathcal{T}|S_1 = i] = 1 + \sum_{j=1}^{10} p_{ij}\tau_j, \tag{10}$$

where $p_{ij}$ is the probability of transitioning from state $i$ to state $j$ (dependent on the stationary policy), and we have already exploited that the goal state does not contribute in the sum because $\tau_G$ is by definition zero.

For a random policy (0.25 probability for each of the four actions) and the layout of the environment in Fig. 1 of the main text, we find $\tau_{\text{trap}} := \tau_8 = \tau_9 = \tau_{10} = \tau_1 + 4$, because it takes on average 4 actions to leave the trap states. Similarly, from state 7, you have a probability of 1/4 to reach the goal in one step, but you can also remain in state 7 or go to one of the trap states. Evaluating all possibilities we arrive at

$$\begin{aligned}
\tau_{\text{trap}} &= \tau_1 + 4, \\
\tau_{i+1} &= 3\tau_i - 2\tau_{\text{trap}} - 4 \;\text{ for }\; i \in \{1, ..., 6\}, \\
\tau_7 &= \frac{4}{3} + \frac{2}{3}\tau_{\text{trap}}.
\end{aligned} \tag{S32}$$

By solving this set of linear equations, we find

$$\begin{aligned}
&\tau_1 = 13116 \;,\; \tau_2 = 13104 \;,\; \tau_3 = 13068 \;,\; \tau_4 = 12960 \\
&\tau_5 = 12636 \;,\; \tau_6 = 11664 \;,\; \tau_7 = 8748 \\
&\tau_8 = \tau_9 = \tau_{10} = \tau_{\text{trap}} = 13120.
\end{aligned} \tag{S33}$$

The results of calculation show that, starting from state 6 (which is the starting state of the first episode in our experiments), it takes on average more than 10000 actions to find the goal with a random policy.

### 6 Precise statement of the prediction in 'Discussion'

Consider an extended version of our environment in Fig. S2 which includes a new (and not necessarily finite) set of states (i.e., the purple states in Fig. S2) that can be accessed from state 4 in the middle of the direct path to the goal. Assume that a participant has found the goal state G at the end of the first episode. In episodes 2 to 5 two different situations may arise. (i) If participants believe that the yellow goal in Fig. S2 is the only (or the most) rewarding state in the environment, then they should ideally stop exploration as soon as they have found the goal and go straight to the goal in subsequent episodes. (ii) If participants wonder whether there may exist another state with a higher value of reward than state G, then they will spend a large amount of time in novelty-rich states like the purple states in Fig. S2. Our prediction, based on the SurNoR model presented

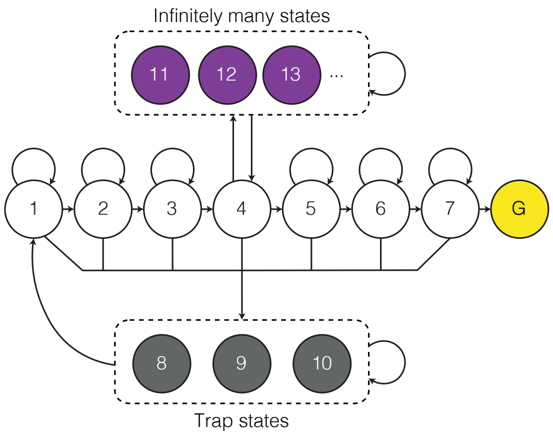


Figure S2: An example of the extended version of our environment mentioned in the 'Discussion' section of the main text. The existence of a set novelty-rich states may distract participants from exploiting the reward at the goal in the episodes after the 1st episode.

in the main text, is that both situations can be observed in the behavioral data and that the difference depends on the prior knowledge given to the participant about the environment before the start of the experiment.

The environment of Fig. S2 also provides a critical test for alternative algorithms of SurNoR. Importantly, in the SurNoR model, information on novelty and external reward are summarized in two separate set of $Q$-values. Consider an alternative model where the novelty is treated as an internal reward and is added to the external reward in a *single* set of $Q$-values. This is equivalent to adding $Q$-values of novelty and reward with a *fixed* factor $\beta_N$. Since the novelty of the purple states is constantly increasing, no matter the values of $\beta$ and $\omega$ in Eq. S24, any fixed and non-zero value of $\beta_N$ (c.f. Eq. S16 and Eq. S18) will eventually drive the agent back towards novelty-seeking and hence exploration of the purple states. This statement holds for non-deterministic model-free, model-based, or hybrid models.

A straightforward way to avoid being distracted from exploiting the external reward is to stop seeking novelty after finding the goal for the first time. This is done in the SurNoR algorithm by reducing $\beta_N$, i.e., by reducing the relative importance of novelty.

### 7 Fitted Parameters

The optimal parameters after fitting the SurNoR model to behavior are summarized in table 2. The reported error for each parameter is the maximum of its standard deviation approximated by Laplace approximation [31] and the optimization precision. Laplace approximation was done for each dimension separately, i.e. the covariance matrix was assumed to be diagonal to avoid the problems arising from approximation of the full Hessian matrix in high-dimensional spaces. Therefore, the reported errors can be seen as lower bounds for the real errors.

| Param. | Value | Error |
|---|---|---|
| $\epsilon$ | 2e−4 | 2e−4 |
| $m$ | 0.31 | 0.02 |
| $\lambda_R$ | 0.97 | 0.01 |
| $\lambda_N$ | 0.70 | 0.05 |
| $\beta_1$ | 4.7 | 0.2 |
| $\beta_2$ | 1.50 | 0.06 |
| $\beta_{N1}$ | 0.145 | 0.006 |
| $\beta_{N2}$ | 0.220 | 0.015 |
| $T_{\text{PS}}$ | 10 | 1 |
| $\mu_R$ | 0.94 | 0.01 |
| $\mu_N$ | 0.80 | 0.05 |
| $Q_{N0}$ | 0.50 | 0.35 |
| $\rho_b$ | 0.06 | 0.02 |
| $\delta\rho$ | 0.45 | 0.05 |
| $\omega_{\text{scale}}$ | 5.8 | 0.2 |
| $\omega_0$ | 0.75 | 0.05 |
| $\omega_{11}$ | 0.20 | 0.05 |
| $\omega_{12}$ | 0.15 | 0.05 |

Table 2: SurNoR parameters fitted to the behavioral data of all participants. This set of parameters was used for EEG analysis and illustrations in Fig. 5 of the main text.
